# Clonal amplification-enhanced gene expression for cell-free directed evolution

**DOI:** 10.1101/2022.11.17.516858

**Authors:** Zhanar Abil, Ana Maria Restrepo Sierra, Christophe Danelon

## Abstract

In cell-free gene expression, low input DNA concentration severely limits the phenotypic output, which may impair in vitro protein evolution efforts. We address this challenge by developing CADGE, a strategy that is based on clonal isothermal amplification of a linear gene-encoding dsDNA template by the minimal Φ29 replication machinery and in situ transcription-translation. We demonstrate the utility of CADGE in bulk and in clonal liposome microcompartments to boost up the phenotypic output of soluble and membrane-associated proteins, as well as to facilitate the recovery of encapsulated DNA. Moreover, we report that CADGE enables the enrichment of a DNA variant from a mock gene library either via a positive feedback loop-based selection or high-throughput screening. This new biological tool can be implemented for cell-free protein engineering and the construction of a synthetic cell.

## Introduction

Directed evolution has revolutionized synthetic biology. Inspired by natural selection, this engineering approach encompasses cycles of genetic diversification and enrichment of rare desired variants, allowing for accelerated protein evolution even with limited a priori knowledge about the structure-function relationships (Romero and Arnold, 2009; Packer and Liu, 2015; Wang et al., 2021). Directed evolution enabled engineering of a plethora of proteins, genetic pathways, and even genomes to generate variants with improved or tailor-made properties (Chatterjee and Yuan, 2006; Johannes and Zhao, 2006; Hida et al., 2007; Turner, 2009; Xiong et al., 2012; Bornscheuer et al., 2019; Engqvist and Rabe, 2019; Zhou et al., 2021). Incorporation of directed evolution principles to the construction framework of a synthetic cell has recently been proposed (Abil and Danelon, 2020). Compartmentalized gene expression in liposomes (Yu et al., 2001; Nomura et al., 2003; Noireaux and Libchaber, 2004) has gained considerable momentum in the last few years, with methodological advances that have improved the yield of functional vesicles (Fujii et al., 2014; Blanken et al., 2019), enabling the reconstitution of complex biological functions, such as DNA replication (van Nies et al., 2018), phospholipid synthesis (Blanken et al., 2020), membrane deformation processes (Godino et al., 2019, 2020; Kattan et al., 2021) and light-triggered ATP synthesis (Berhanu et al., 2019). Moving forward to optimizing and integrating cellular modules may require a system’s level evolutionary approach (Abil and Danelon, 2020).

Over the past decades, numerous in vivo and cell-free methodologies for gene expression of the targeted phenotypes have been developed. In vivo methodologies have been the most common, since a suitable host organism could provide low-cost gene expression with a reliable yield (Pourmir and Johannes, 2012). However, cell-free systems have emerged as an alternative and attractive platform due to the higher degree of controllability and freedom from the constrains related to cell survival (Shimizu et al., 2006; Perez et al., 2016). Cell-free protein synthesis enabled engineering of a number of proteins, including membrane or cytotoxic proteins (Fujii et al., 2015; Holstein et al., 2021) as well as peptides and proteins that incorporate unnatural amino acids (Watts and Forster, 2012; Uyeda et al., 2015; Newton et al., 2020). Cell-free protein expression can be accomplished using either cell lysates (Silverman et al., 2020) or in a reconstituted transcription-translation system such as the PURE system (Shimizu et al., 2001).

A pre-requisite for directed evolution is a genotype-phenotype link. In cell-free systems, this link is often implemented through ribosomal, mRNA, or other cell-free macromolecular display technologies (Dodevski et al., 2015), although these techniques are often limited to evolution of peptide and protein binding affinities. For evolution of an enzyme’s catalytic turnover, however, compartmentalization in emulsion droplets (Tawfik and Griffiths, 1998) or liposomes (Sunami et al., 2006) is more appropriate. Such biomimetic compartments are also often used as the chassis for engineering towards construction of an artificial cell (Cho and Lu, 2020). Finally, liposomes are exceptionally suited for evolution of membrane proteins requiring a lipid bilayer for solubility and activity (Fujii et al., 2015).

However, coupling the gene expression and enrichment steps in a cell-free system within a microcompartment is often limited by the low yield of synthetized proteins from a single DNA template. Although detectable activity of cell-free expressed proteins arising from a single gene copy has been demonstrated in some experimental conditions (Nishikawa et al., 2012; Fujii et al., 2013; Uyeda et al., 2016; Zhang et al., 2019), it is hardly surprising that below a certain threshold, template DNA concentration is a limiting factor for in vitro protein expression (Chory and Kaplan, 1982; Noireaux and Libchaber, 2004; Stögbauer et al., 2012; Niederholtmeyer et al., 2013; Nourian and Danelon, 2013). In fact, production of full-length proteins in reconstituted systems ceases before NTPs and amino acids get depleted, and efforts to increase the amount of protein from low DNA concentrations remain frustrated (Doerr et al., 2019). Therefore, clonal amplification of expression templates is a generic solution to enhance protein yield and activity readout, as well as the recovery of selected DNA variants.

Several in vitro methods based on polymerase chain reaction (PCR) or isothermal amplification have been utilized for clonally amplifying DNA templates prior to gene expression and enrichment of variants. In particular, rolling circle amplification (RCA) based on the Φ29 DNA polymerase and replication cycle reaction (RCR) based on a reconstituted *Escherichia coli* replisome are compatible with droplet microcompartments (Holstein et al., 2021; Ueno et al., 2021). While RCR has not been combined with gene expression yet, RCA and PCR methods are not fully compatible with cell-free protein synthesis due to cross-inhibition by some components or different incubation temperatures. Therefore, multiple-step workflows have been implemented, which require droplet-based microfluidic handling (Mazutis et al., 2009; Fallah-Araghi et al., 2012; Galinis et al., 2016; Holstein et al., 2021) or bead-display (Diamante et al., 2013; Paul et al., 2013; Plesa et al., 2018; Lindenburg et al., 2020; Restrepo Sierra et al., 2022). Still, a major challenge in cell-free directed evolution is the coupling of DNA amplification from single template copies, gene expression and quantitation of the activity of the protein of interest for fitness assignment in *one* environment.

In this study, we simplify the in vitro evolution methodology by a single isothermal, <u>c</u>lonal <u>a</u>mplification-enhance<u>d g</u>ene <u>e</u>xpression, or CADGE. The strategy relies on the protein-primed replication machinery of the *Bacillus subtilis* bacteriophage Φ29 (Mencía et al., 2011) consisting of DNA polymerase (DNAP, encoded by gene *p2*), terminal protein (TP, encoded by gene *p3*), double-stranded DNA-binding protein (DSB, encoded by gene *p6*), and single-stranded DNA-binding protein (SSB, encoded by gene *p5*), and requires prior flanking of the gene of interest (GOI) with Φ29 origins of replication (ori) using a standard recombinant DNA technique of choice. The Φ29 DNAP is chosen largely due to its strand-displacement activity, a relatively rare property for a family B DNA polymerase (Kamtekar et al., 2004; Rodríguez et al., 2005). This activity enables it to displace the non-template DNA strand at ambient temperatures, thus ensuring compatibility with cell-free transcription-translation. In addition, Φ29 DNAP has an excellent processivity (Blanco et al., 1989; Rodríguez et al., 2005), which could be useful at efficiently replicating long and multigene DNA templates. To initiate the replication, DNAP forms a complex with TP (Kamtekar et al., 2006), and the heterodimer is recruited to replication origins, a process that is facilitated by DSB (Blanco et al., 1986). DSB activates the replication initiation by forming a multimeric nucleoprotein complex at the origins of replication (Freire et al., 1996), whereas TP primes the DNA synthesis at each end, remaining covalently attached to the 5’-end of the daughter strand (Penalva and Salas, 1982). After successful priming, DNAP dissociates from the complex and continues the polymerization activity (Méndez et al., 1997). SSB is another auxiliary protein, which assists in the replication by stabilizing the displaced strand (Martìin et al., 1989). Using this system, we previously realized transcription-translation-coupled self-replication of a two-gene construct (van Nies et al., 2018). Herein, we demonstrate that transcription-translation-coupled amplification of orthogonal genes can be achieved in bulk and in liposome compartments, improving the expression level of a GOI. As a proof-of-concept, we show the enrichment of an ori-*GOI* from a mock library encapsulated in liposomes, a key step toward cell-free protein evolution. Moreover, we apply CADGE to enable the screening of protein functions that are relevant in the field of synthetic cell construction.

## Results

### Design of CADGE

The CADGE strategy involves the following minimal requirements (Fig. 1a,b):

**Figure 1.**
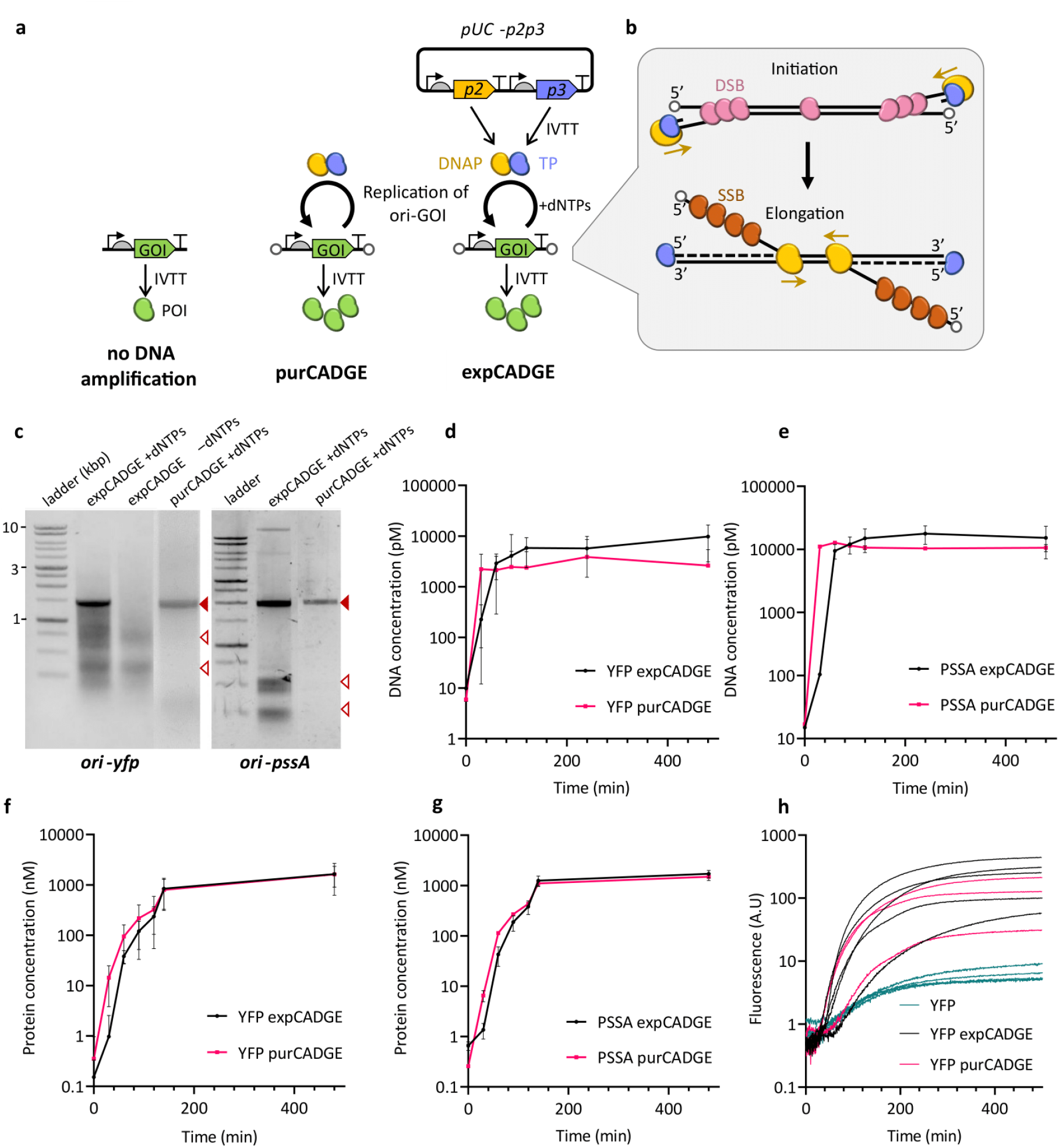
Principles and validation of the CADGE strategy in bulk reactions. **(a)** Schematics of the different gene expression configurations used in this study. **(b)** Schematic of linear DNA replication by the Φ29 minimal DNA replication machinery. **(c)** Visualization of amplified DNA in CADGE samples via agarose gel electrophoresis. Filled red arrowheads indicate expected product size, empty red arrowheads indicate unfinished smaller products. **(d, e)** Time-course analysis of ori-*yfp* (d) and ori-*pssA* (e) DNA amplification in bulk CADGE reactions via absolute qPCR quantitation. **(f**,**g)** Time-course analysis of YFP and PssA expression in bulk CADGE reactions via absolute LC-MS/MS quantitation. **(h)** Time-course analysis of relative YFP expression in bulk CADGE reactions via fluorescence measurements in a microplate reader. Error bars indicate standard deviation from three (data sets in d, e, f) to four (data set in g) biological replicates.

1. A GOI is inserted between the 191-bp-oriL and 194-bp-oriR origins of replication of the Φ29 genome, although the 68-bp minimal origins could potentially be used as well (Serrano et al., 1989). The DNA template must be linearized with the origins at each end of the molecule, which can be achieved by PCR amplification from ori-containing plasmid DNA. Moreover, the linear DNA has to be phosphorylated at each 5’-end, which can be done by using 5’-phosphorylated primers. One, two (van Nies et al., 2018) or, in principle, more genes can be encoded on a single ori-flanked DNA template. Hereafter, we refer to such linear constructs as ori-*GOI*.
2. The PURE*frex2*.*0* system is chosen for in vitro transcription-translation because of its higher purity and reduced nuclease activity compared to other commercial PURE systems (Doerr et al., 2019). Thus, the linear DNA construct contains regulatory elements compatible with gene expression in PURE*frex2*.*0*. These comprise a T7 promoter, g10 leader sequence, *E. coli* ribosome binding site and a transcription terminator (e.g. T7 and vesicular stomatitis virus terminators).
3. The system requires four minimal protein components of the Φ29 replication machinery: DNAP, TP, SSB, and DSB, plus dNTPs and ammonium sulphate for the efficient dimerization of the replication initiation complex (Blanco et al., 1987) (Fig. 1a,b). DNAP and TP can either be introduced in a purified form (purCADGE) or in situ expressed from a separate DNA construct (expCADGE). In the latter configuration, the two genes *p2* and *p3* are introduced on a single plasmid, self-replication being prohibited by the circular nature of the DNA. Although SSB and DSB can be functionally expressed in the PURE system (Van Nies et al., 2018), we recommend supplying them as purified proteins since they are required at micromolar concentrations and their cell-free expression would create a burden on the transcription-translation apparatus. The linear replication product in CADGE is essentially identical to the parental DNA molecule – except for the fact that TP is covalently bound at the 5’-end of each daughter strand. In the current protocol, the 5’-TP is lost with subsequent PCR amplification during recovery of the total DNA from liposomes. Thus, the resulting recovered DNA is identical in its structure to the original template DNA, and does not require any additional processing between rounds of evolution.

### Validation of CADGE in bulk reactions

We first evaluated the performance of CADGE on amplifying a GOI in bulk reactions. Two ori-GOI fragments encoding either the enhanced yellow fluorescent protein (eYFP) or *Escherichia coli* phosphatidylserine synthase (PssA) (ori-*yfp* and ori-*pssA*, respectively) were constructed. The DNA template was added to PURE*frex2*.*0* in the presence of the DNA replication machinery, dNTPs, and ammonium sulphate, and the solution was incubated at 30 °C. With both purCADGE and expCADGE, we found that the full-length DNA (Fig. 1c) can be amplified to saturation by two to three orders of magnitude from an input template concentration of 10 pM within two hours, as confirmed by absolute quantification by qPCR (Fig. 1d,e). Although full-length replication products are the most abundant DNA species, shorter fragments are also visible in the gel, especially with expCADGE (Fig. 1c).

To test if template amplification is accompanied with an increase in protein expression levels, we quantified the concentrations of eYFP and PssA by liquid chromatography-coupled mass spectrometry (LC-MS), and confirmed the production of both proteins to up to 1 μM, with no noticeable differences between purCADGE and expCADGE (Fig. 1f,g). These amounts of protein expression were comparable to the generally reported cell-free protein synthesis levels (Carlson et al., 2012)(Godino et al., 2019)(Doerr et al., 2021), but with considerably (two to three orders of magnitude) less input of template DNA. Importantly, fluorescence kinetics measurements show that in the absence of DNA amplification, only a very low level of YFP was expressed even after several hours of incubation (Fig. 1h). This finding indicates that the enhanced protein expression is the direct result of gene amplification. Kinetic analysis of protein synthesis gives apparent maximum translation rates (defined as the highest slope) and time before saturation of about 1.2 nM min^−1^ and 300 min, respectively. These values are consistent with previous data obtained with nanomolar concentrations of DNA template (Blanken et al., 2019)(Doerr et al., 2019), suggesting that CADGE does not significantly delay or slow down protein production.

### CADGE improves phenotypic output in liposomes

We next sought to demonstrate that CADGE is able to increase the number of liposomes with detectable amounts of synthesized protein starting from clonal quantities of DNA molecules (Fig. 2a). To this end, the construct ori-*yfp* and the CADGE components were encapsulated in a polydisperse population of liposomes, the bilayer of which is composed of biologically relevant lipids found in the composition of *E. coli* plasma membrane (van Nies et al., 2018). Ori-*yfp* was introduced at 10 pM bulk concentration, corresponding to an expected average number of DNA molecules per liposome *λ* = 0.2 (Methods section) if one assumes an average liposome radius of 2 μm (Blanken et al., 2020). To confine the in vitro transcription-translation (IVTT) and replication reaction to the interior of the liposomes, we introduced DNase I to the outer phase of the liposome population, which resulted in further reduction of the overall input DNA concentration to around 100 fM (Fig. 2b). The extent of clonal amplification was assessed by comparing endpoint data (typically overnight incubation at 30 °C) with (+) and without (–) dNTPs. To recover the DNA for analysis, we heat-inactivated DNase I and released the DNA from the liposomes by dilution in water. Quantification by qPCR revealed that in both purCADGE and expCADGE, over 100-times more DNA was obtained in the presence of dNTPs than in the absence thereof (Fig. 2b), suggesting that DNA was amplified inside the liposomes.

**Figure 2.**
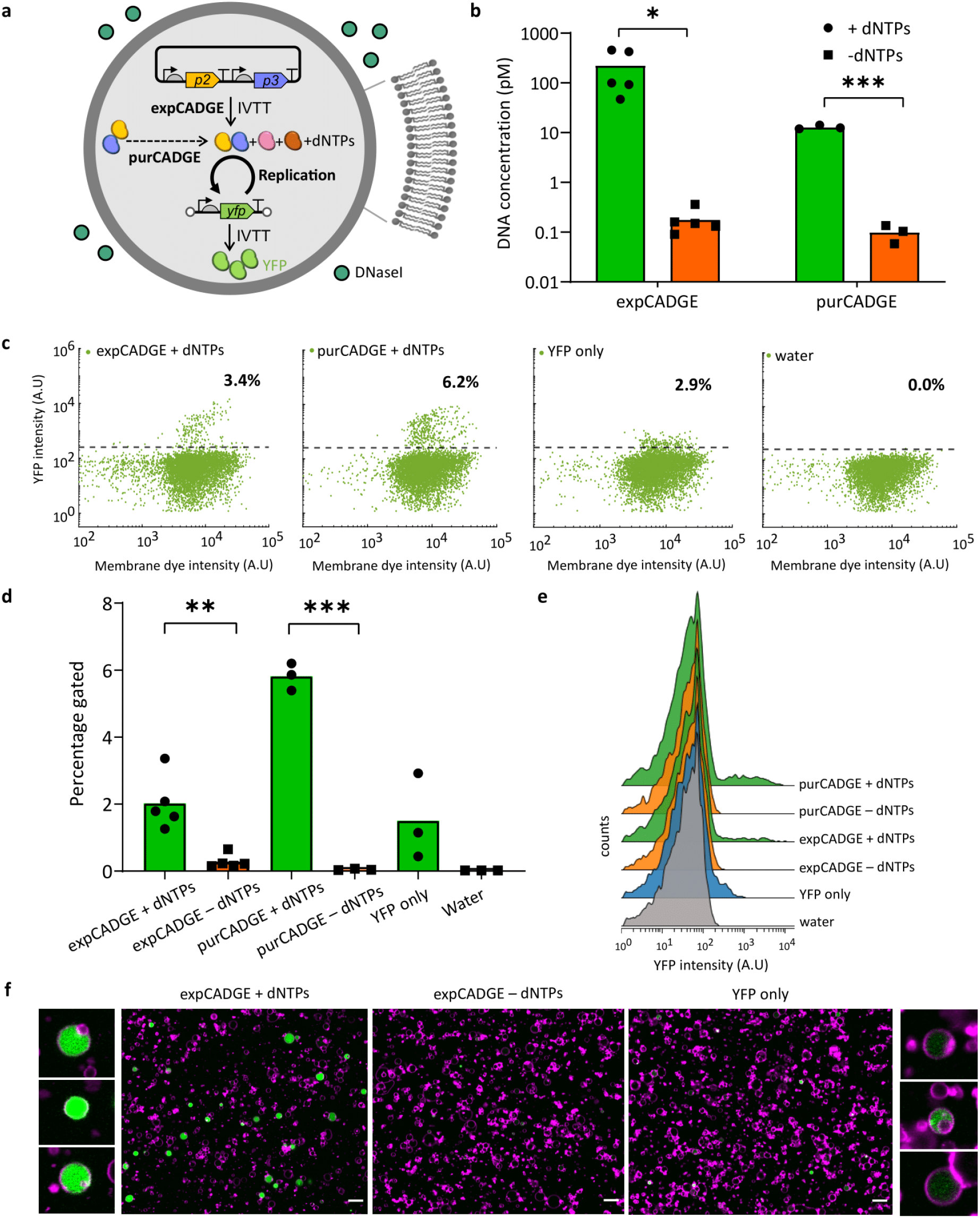
Effect of in-liposome CADGE on the phenotypic output of a reporter gene. **(a)** Schematic of in-vesiculo reporter gene amplification and expression via CADGE. **(b)** Absolute quantitation of ori-*yfp* DNA by qPCR in lysed liposomes. Total 10 pM input template DNA concentration was used, which was reduced due to externally supplied DNaseI. **(c)** Populational variation of in-vesiculo YFP fluorescence in CADGE samples measured by flow cytometry. **(d)** Quantitation of the fraction of liposomes showing above-background YFP fluorescence estimated by the horizontal gate in (c). **(e)** Histogram representation of flow cytometric data of YFP expression in CADGE liposomes. **(f)** Confocal microscopy imaging of CADGE and un-amplified samples. Magenta: Texas Red (membrane dye). Green: YFP. Scale bars: 5 μm. *P ≤ 0.05; ** P ≤ 0.01; *** P ≤ 0.001.

To assess the effect of DNA amplification on gene expression, we analyzed individual liposomes for YFP signal using flow cytometry (Fig. 2c,d,e). We confirmed that under both CADGE configurations, and in the presence of dNTPs, a higher percentage of liposomes exhibited a YFP fluorescence above the background level (Fig 2c,d). This was expected from the strongly reduced protein expression level at low DNA concentrations (Fig. S1). Interestingly, the intensities of YFP-expressing liposomes were higher in the presence of dNTPs compared to those in the absence but also compared to samples that contained none of the components of the minimal replication machinery (Fig. 2c,e,f). Similar observation was made from fluorescence imaging of individual liposomes (Fig. 2f). These results suggest that clonal amplification does not only boost gene expression to overpass the threshold for measurable activity, but also increases the dynamic range of the phenotypic output. Although the percentage of YFP-expressing liposomes was slightly higher with purCADGE compared to expCADGE (Fig. 2d), the intensity profiles were similar (Fig. 2e), suggesting that co-expression of *p2* and *p3* genes does not significantly affect the production of protein of interest (POI) in liposomes. Similar conclusion could be reached from bulk reactions (Fig. 1f,g).

We noticed that the percentage of YFP-expressing liposomes was lower in the –dNTPs sample when compared to the condition where replication reagents were omitted (YFP only, Fig. 2d). This suggests that some replication components may inhibit transcription-translation. We tested this hypothesis by varying DSB and SSB concentrations in ori-*yfp* bulk reactions and found that reduced amounts of DSB led to higher expression of ori-*yfp*, while changing SSB concentrations had little effect (Fig. S2). Considering that DSB is a Φ29 transcription regulator of early and late genes (Meijer et al., 2001), it is possible that binding of DSB to the DNA template inhibits gene expression in vitro. Therefore, we decided to lower DSB concentration down to either 52.5 or 105 μg/mL, in order to mitigate inhibitory effects without compromising DNA replication.

### CADGE with a positive feedback loop

We then implemented expCADGE with a positive feedback loop coupling POI synthesis back to DNA replication. The autocatalytic framework of this selection strategy may offer a more effective and efficient alternative to fluorescence-based screening methods. Ori-*p3* template coding for TP was introduced at 10 pM concentration (*λ* = 0.2), supplemented with an excess amount of plasmid encoding solely the DNAP (Fig. 3a), and encapsulated in liposomes. We hypothesized that an initial seed expression of TP could kick off the replication of ori-*p3* with the expressed DNAP and yield increasing amounts of ori-*p3*. Quantitative PCR showed that the *p3* gene was amplified inside liposomes by three orders of magnitude in the presence of dNTPs (Fig. 3b) compared to the –dNTPs control. The DNA intercalating dye dsGreen was used as a fluorescent marker to assess DNA amplification in single vesicles by flow cytometry. A fraction of liposomes with increased dsGreen fluorescence compared to the background was detected in the presence of dNTPs, which corroborates that self-amplification of DNA took place (Fig. 3c,d).

**Figure 3.**
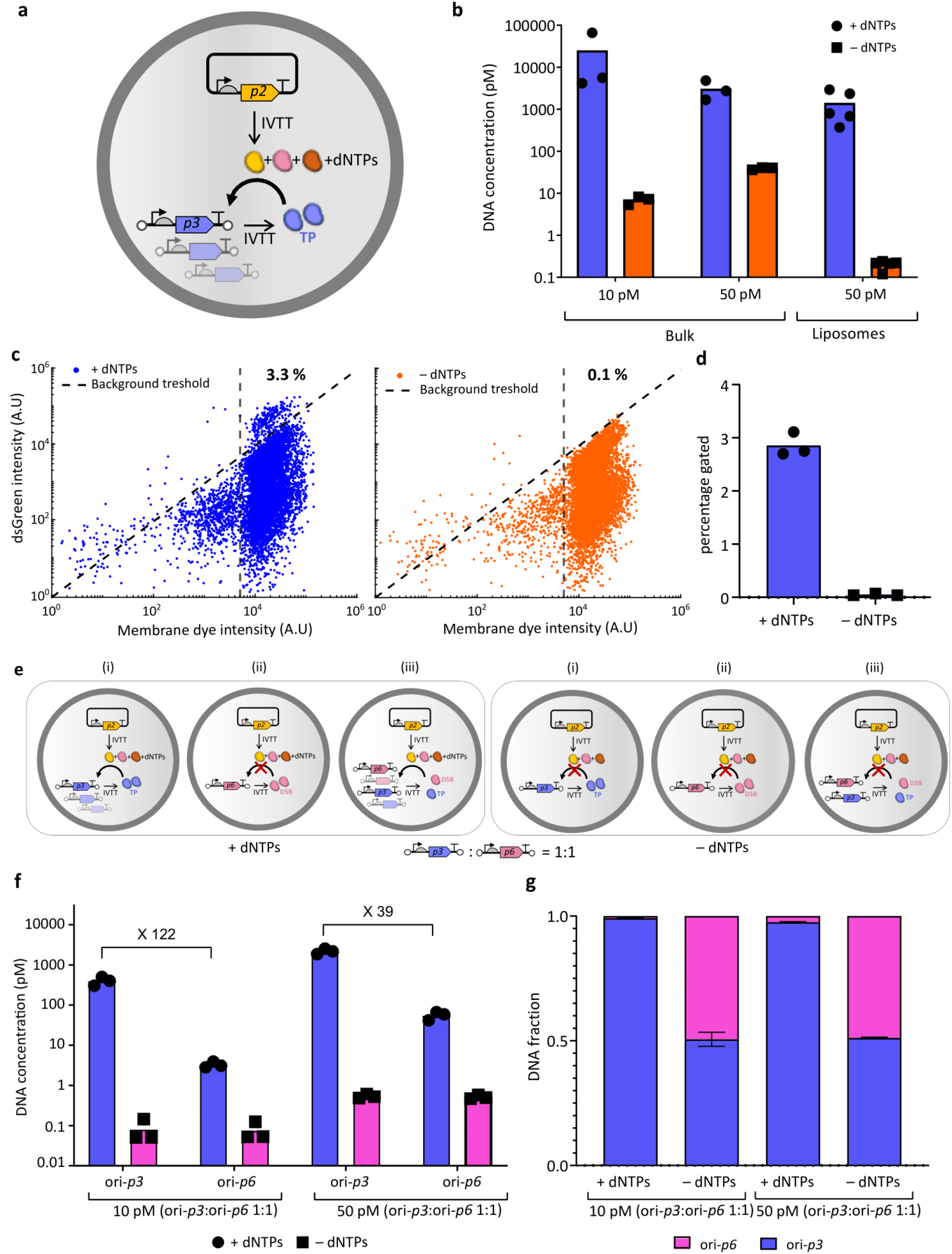
CADGE with a positive feedback loop. **(a)** Schematic of in-vesiculo ori-*p3* DNA amplification and expression via CADGE. **(b)** Absolute quantitation of ori-*p3* DNA by qPCR in lysed liposomes. Total 10 pM input template DNA concentration was used, which was reduced due to externally supplied DNaseI. **(c)** Populational variation of dsGreen fluorescence in CADGE liposomes measured by flow cytometry. **(d)** Quantitation of the fraction of liposomes showing above-background dsGreen fluorescence estimated by the diagonal gate in (c). **(e)** Schematics of the different gene expression configurations used in (f) and (g). **(f)** Absolute DNA quantitation of two genes by qPCR of liposome suspensions after a single round of selection. Total 10 pM or 50 pM input template DNA concentration was used, which was reduced due to externally supplied DNaseI. **(g)** Calculated fractions of the two genes in the mixture after a single round of selection. Error bars indicate standard deviation from three biological replicates.

The high amplification of ori-*p3* prompted us to experimentally determine the bulk concentration of DNA template below which the amplification is ‘clonal’. Experimental validation of the *λ* = 1 regime is important considering the polydispersity of the liposome population, which differs from our assumption of constant volume (Methods section). Therefore, we performed a mock enrichment experiment by co-encapsulating ori-*p3* and an equimolar amount of unrelated DNA, also flanked with replication origins (here ori-*p6*) (Fig. 3e). In this scenario DNA replication is conditional to the presence of both DNAP and TP. Therefore, ori-*p6* can only be replicated when co-encapsulated with ori-*p3*, i.e. under nonclonal conditions where more than a single molecule of ori-*GOI* is present in a liposome. Conversely, an enrichment of ori-*p3* over ori-*p6* would indicate that amplification is mostly clonal. After a single round of selection, ori-*p3* was enriched 122-fold and 39-fold over ori-*p6* when starting with 10 pM and 50 pM DNA concentrations, respectively (Fig. 3f,g). This result confirms that in-liposome amplification of ori-*GOI* is mostly clonal and that CADGE is suitable for in vitro directed evolution purposes.

Amplification of ori-*p6* was however not totally prohibited, even at 10 pM input mixture concentration (Fig 3f). The latter is not unexpected considering that the estimated probability of co-occupancy of the ori-*p3* and ori-*p6* templates is not zero but is (1 − *e*^−λ^)_2_ = 0.15 with 50 pM input mixture concentration (*λ* = 0.5 for each ori-*GOI*). Together, the significant enrichment of ori-*p3* over ori-*p6* experimentally validates that 10 pM (and to some extent 50 pM) concentration is enough to keep a strong genotype to phenotype link in our polydisperse liposome population. This experiment also implies that, as long as DNA replication can be coupled to a POI activity, TP or any other POI can be potentially evolved using this selection scheme.

### CADGE improves enrichment efficiency of a GOI based on high-throughput screening

Next, we asked whether CADGE may be beneficial for in vitro protein evolution via fluorescence-based screening. To this end, we performed a mock enrichment experiment at a clonal expression condition with 10 pM ori-*GOI*, i.e. *λ* = 0.2. We aimed to enrich the DNA template ori-*yfp* from a mock library containing an excess of the unrelated template ori-*minD* based on the fluorescence of expressed YFP by fluorescence activated cell sorting (FACS). For this, the ori-*yfp* DNA template was mixed with 10-fold excess of ori-*minD* template and the DNA/PURE mixture was encapsulated in liposomes at a total 10 pM DNA concentration (Fig. 4a). At such a low template DNA concentration (1 pM of ori-*yfp*), expression of YFP is significantly reduced compared to higher DNA concentrations typically used in cell-free reactions, leading to low signal-to-noise ratio (Fig 4b). As expected from previous results, CADGE liposomes exhibited higher dynamic range of YFP fluorescence compared to liposomes that contained the same input DNA mixture concentration but no replication factors (Fig. 4b,c). For sorting, two stringency conditions were tested: the “all-gate”, which encompassed the top 1% of all the liposomes (applied to both non-amplified and CADGE samples), and the “high-gate”, which included only the top (0.2%) of the high-intensity liposomes (applied to CADGE samples only). It was reproducibly difficult to recover the full-length DNA by PCR from the non-amplified liposome samples, while full-length DNA from liposomes with implemented CADGE was easily recovered (Fig. 4e). This finding can be explained by higher DNA titers in the sorted liposomes from CADGE samples. Indeed, as assayed by qPCR, ori-*yfp* and ori-*minD* mixtures in liposomes were comparably (both more than a 100-fold) and uniformly (i.e. two genes amplified equally in a single sample) amplified with both purCADGE and expCADGE (Fig 4f-h pre-sorted samples). Furthermore, qPCR quantification of sorted liposome samples suggests that using the more stringent condition of “high-gate” in CADGE samples results in improved purity of YFP sorting compared to “all-gate” in both CADGE samples (Fig. 4h; Fig. S3). These findings show that CADGE improves enrichment efficiency and DNA recovery in a single round of mock enrichment experiment, and thus suggest that CADGE may facilitate in vitro protein evolution.

**Figure 4.**
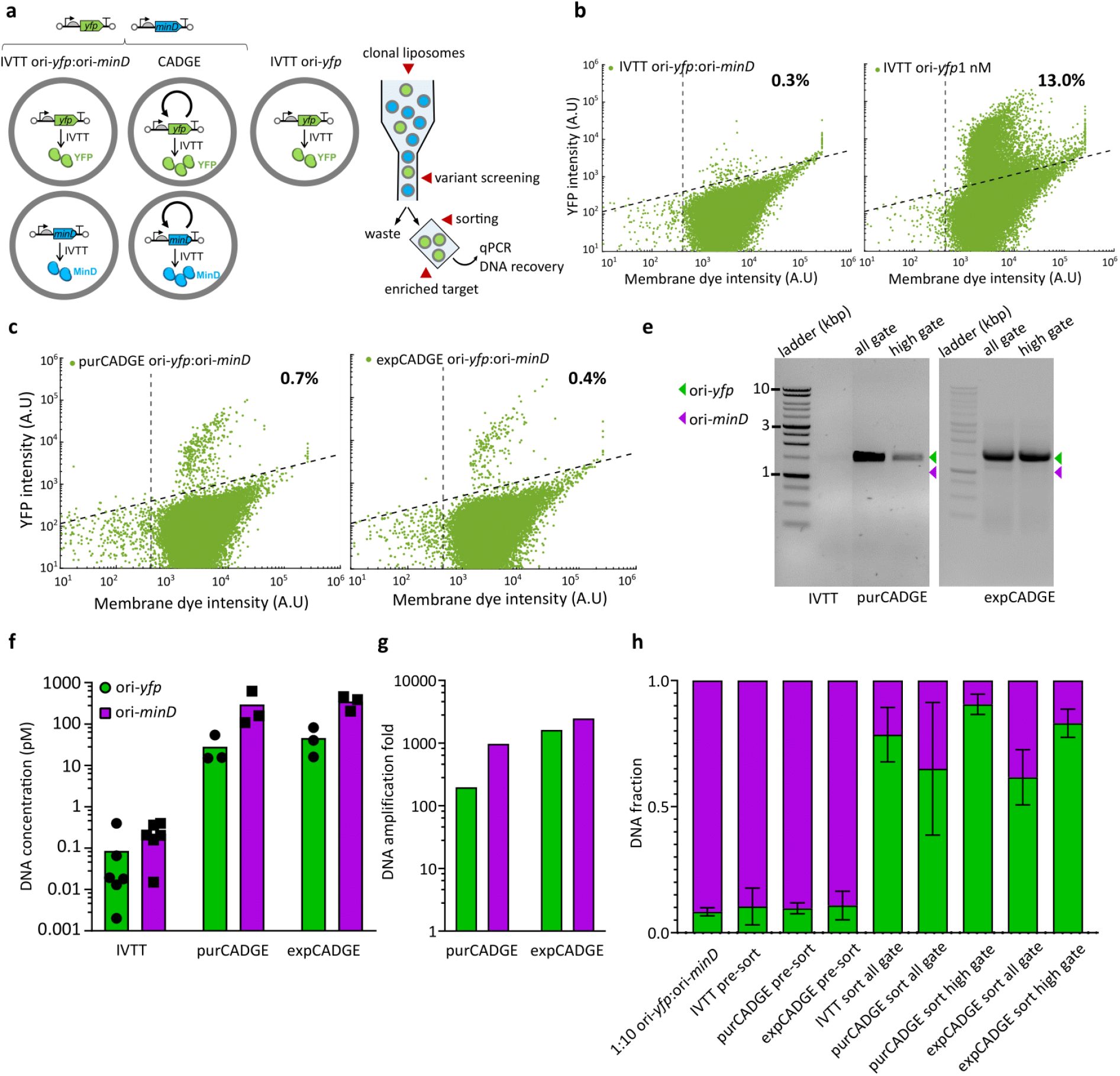
Enrichment of clonally amplified GOI via high-throughput FACS screening. **(a)** Schematics of clonal gene expression and enrichment experimental design for FACS. **(b)** Flow cytometry analysis of liposomes prepared from indicated DNA template mixtures in the PURE system: 10 pM total with 1:1 YFP:MinD DNA mixture (left) and 1 nM YFP DNA (right). **(c)** Flow cytometry analysis of CADGE liposomes prepared from 10 pM total input YFP:MinD DNA template mixture. **(e)** Agarose gel electrophoresis images of recovered DNA from sorted CADGE liposomes. Green arrowhead indicates the expected DNA size for ori-*yfp* (2 kb), purple arrowhead indicates the expected size for ori-*minD* (∼ 1.4 kb). **(f)** Absolute DNA quantitation by qPCR of pre-sort liposome suspensions. **(g)** DNA amplification in pre-sort purCADGE and expCADGE liposome samples. Amplification was calculated as DNA concentration of a specific gene in end-point samples with dNTPs compared to end-point samples without dNTPs. Color coding is the same as in (f). **(h)** Fractions of ori-*minD* and ori-*yfp* DNA mixtures recovered from pre-sort or post-sort liposomes as calculated from absolute DNA quantification by qPCR. Error bars indicate standard deviation from three to seven biological replicates.

### CADGE improves phenotypic output of synthetic cell modules

We previously proposed in vitro evolution as a route to build a synthetic minimal cell (Abil and Danelon, 2020). Here, we seek to exploit CADGE for improving the expression of genes that are relevant for the construction of functional cellular modules. One candidate gene is *pssA* from the Kennedy phospholipid biosynthesis pathway (Raetz, 1976; DeChavigny et al., 1991). The *E. coli pssA* gene encodes an enzyme that conjugates cytidine diphosphate-diacylglycerol (CDP-DAG) with L-serine to produce cytidine monophosphate and phosphatydilserine (PS) (Fig. 5a), a precursor of phosphatidylethanolamine. To assay the activity of in vesiculo synthesized PssA enzyme, we encapsulated PURE and the plasmid encoding the *pssA* gene in phospholipid liposomes containing 5 mol% CDP-DAG, and digested the extraliposomal DNA with DNaseI (Fig. 5b). Since PssA is active as a membrane-associated enzyme, PS would be incorporated on the inner leaflet of the membrane (Hirabayashi et al., 1976; Larson and Dowhan, 1976). However, as previously suggested (Blanken et al., 2020), we expected some flipping of phospholipids to the outer membrane, such that the enzymatic activity of entrapped PssA could be detected by externally staining the liposomes with a PS-specific probe. To this end, we implemented C2-domain of lactadherin protein (LactC2) fused to a fluorescent protein like mCherry or eGFP (Fig. 5b) (Blanken et al., 2020). By flow cytometric analysis of LactC2-mCherry- and Acridine Orange-(membrane marker) stained liposomes, we observed that PS production (and, we assumed, gene expression) reduces considerably at limiting template DNA concentrations (10 and 50 pM DNA) compared to 1 nM (Fig. 5c,d; Fig. S4). Alternatively, we stained the liposomes with LactC2-eGFP and Texas Red (membrane dye), and imaged them by confocal microscopy (Fig. S5). We found that limiting the template DNA concentration visibly reduces LactC2 binding, suggesting that *pssA* gene expression is diminished at low input DNA concentrations.

**Figure 5.**
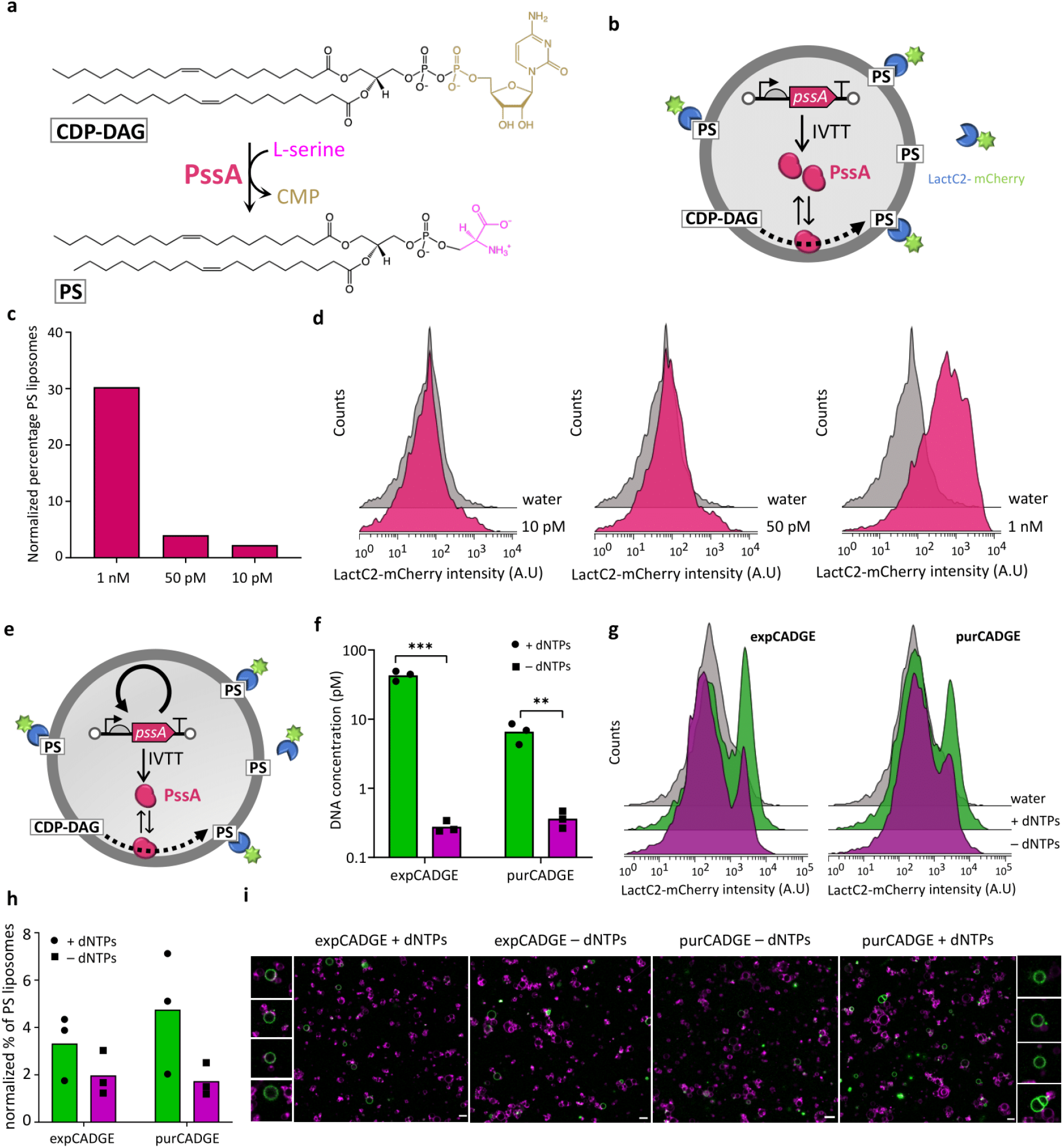
Application of CADGE for improving the enzymatic catalysis of phospholipid headgroup conversion. **(a)** Schematic of CDP-DAG conversion to PS by PssA. **(b)** Schematic of in-vesiculo-expressed PssA enzyme activity and detection of PS-positive liposomes by LactC2-mCherry binding. Percentage quantitation **(c)** and histograms **(d)** of PS-positive liposomes expressing PssA in the PURE system, as assayed by flow cytometry (raw data in Fig S4, S5). **(e)** Schematic of CADGE liposomes expressing PssA enzyme and detection of PS-positive liposomes by LactC2-mCherry binding. **(f)** Absolute DNA quantitation by qPCR of lyzed CADGE liposome samples. ** P ≤ 0.01; *** P ≤ 0.001. **(g)** Histograms and **(h)** quantitation of PS-positive CADGE liposomes expressing PssA, as assayed by flow cytometry (raw data in Fig S6, S7). **(i)** Confocal microscopy of CADGE liposomes expressing PssA. Green, LactC2-eGFP; Magenta, Texas Red-conjugated lipids. Scale bars: 5 μm.

To test if clonal DNA amplification can improve PssA expression, we co-encapsulated the linear ori-*pssA* DNA fragment (10 or 50 pM) with the required additives for either purCADGE or expCADGE (Fig. 5e) and incubated at 30 °C for 4 hours. Using qPCR, we confirmed 10-to 100-fold amplification of the *pssA* gene compared to –dNTP controls with input ori-*pssA* concentrations of 10 pM, in both CADGE configurations (Fig. 5f). Even though PS synthesis was detectable in –dNTP samples, the number of liposomes exhibiting a PS-positive phenotype increased with functional CADGE (+dNTPs, Fig. 5g,h,i; Fig. S6, S7). Overall, these findings demonstrate that, within a synthetic cell context, clonal amplification of template DNA can improve phospholipid headgroup conversion from in vitro expressed PssA protein.

Besides gene-directed phospholipid production, we decided to explore the benefit of CADGE for implementation of the Min system in clonal liposomes. The Min system is involved in the spatial organization of cytokinesis events in *E. coli* (Rowlett and Margolin, 2015) and is therefore a relevant protein system for synthetic cell division. MinD is an ATP-dependent membrane binding protein that recruits MinC, an FtsZ-polymerization inhibitor (Rowlett and Margolin, 2013). We assembled expCADGE reactions with 10 pM ori-*minD* DNA and purified eGFP-MinC as a reporter of the activity of synthesized MinD (Fig. 6a), and encapsulated the mixture in liposomes. Quantitative PCR data showed that ori-*minD* was clonally amplified almost a thousand-fold compared to –dNTPs control samples (Fig. 6b). Confocal imaging and analysis of eGFP-MinC fluorescence distribution in the lumen and at the membrane revealed that in the vast majority of the liposomes, the basal amount of expressed MinD is not sufficient to recruit eGFP-MinC to the membrane (–dNTPs, Fig. 6c,d,e). Using expCADGE, a larger fraction of liposomes exhibited an excess fluorescence signal at the membrane (+dNTPs, Fig. 6c,d,e), indicating that clonal amplification led to a re-localization of MinC through improved expression of functional MinD.

**Figure 6.**
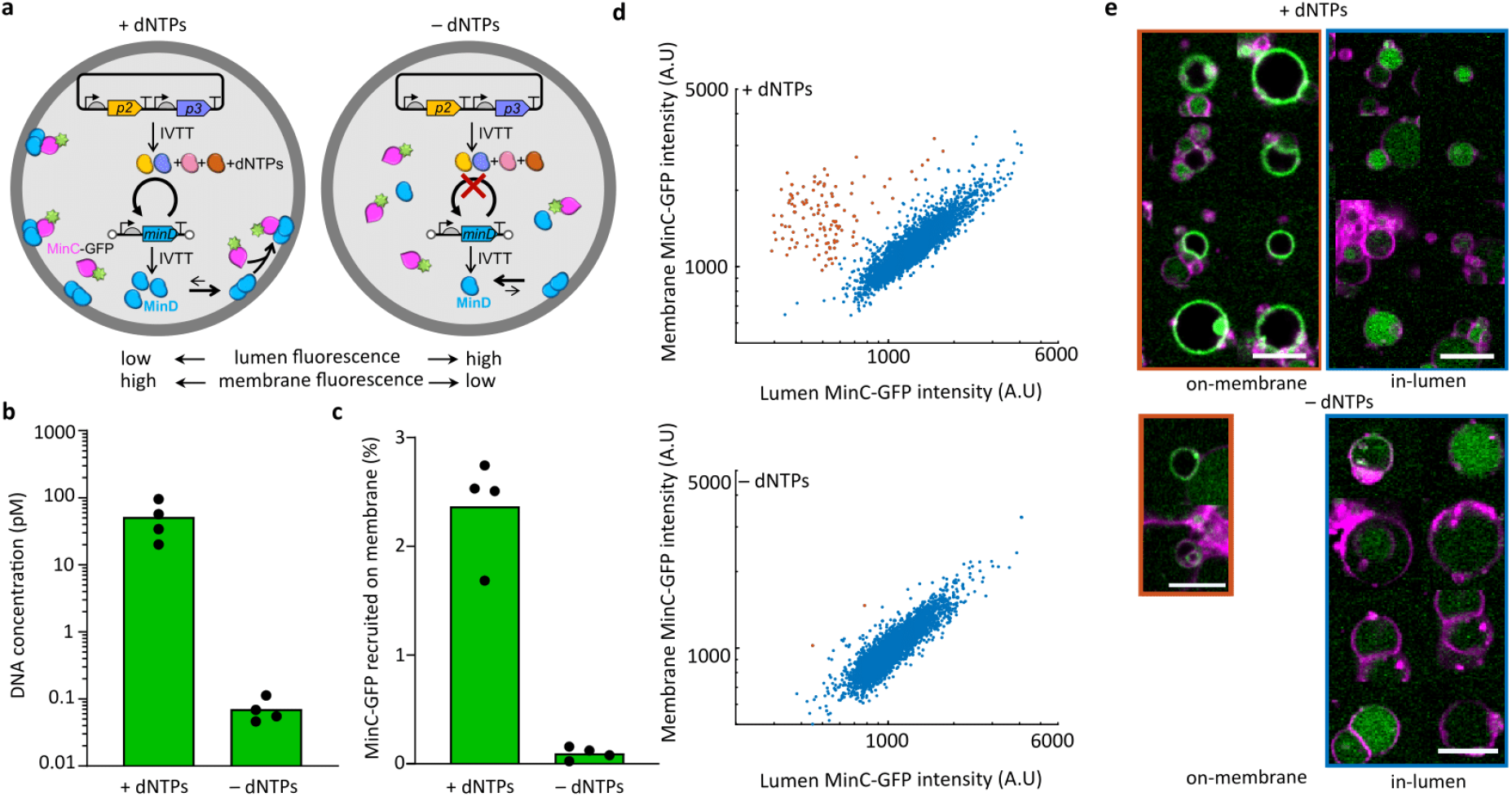
Application of CADGE for facilitating the membrane recruitment of Min proteins. **(a)** Schematics of MinC-MinD membrane binding assay in expCADGE liposomes. **(b)** Absolute quantitation of ori-*minD* DNA by qPCR of lyzed expCADGE liposome samples. Individual symbols are independent biological repeats. **(c)** Percentage of MinC-GFP recruited to the membrane, as obtained from fluorescence confocal images. **(d)** Scatter plots of the membrane and lumen fluorescence intensities in individual liposomes. The liposomes marked in orange dots display a marked recruitment of MinC-GFP to the membrane. **(e)** Montage of randomly selected liposome images taken from the data shown in **(d)**. Scale bars: 5 μm.

## Discussion

Herein, we established CADGE, a single-step DNA amplification and in situ transcription-translation strategy that can be used for improving clonal gene expression. Our findings suggest that CADGE can be instrumental in facilitating in vitro evolution of a variety of genes, including those that are important for synthetic cell construction. The general applicability of this strategy is enabled by only a few requirements: the possibility of in vitro expression of the GOI, and in vitro activity of the POI.

Advantages of CADGE for clonal amplification, compared to previous strategies (Mazutis et al., 2009; Fallah-Araghi et al., 2012; Diamante et al., 2013; Paul et al., 2013; Lu et al., 2014; Galinis et al., 2016; Lindenburg et al., 2020; Holstein et al., 2021) include minimum effort (i.e. a single-step amplification and expression), time (around two hours to set up the reaction), and instrumentation (although microfluidic devices for microcompartmentalization or screening can be implemented, if necessary). The benefit of using CADGE post-enrichment lies in the simplicity of the protocol since only PCR is required to proceed to another round of encapsulation/enrichment, and in the improved DNA recovery yield.

Despite a number of advantages, CADGE is not without some limitations. Thus, DSB was found to exhibit some inhibitory effect on gene expression, which could be mitigated to some extent by optimized DSB concentrations (Fig. S2). Moreover, in our fluorescence measurements of CADGE samples, the measured percentage of liposomes with a fluorescence signal above the activity threshold (3 to 5% with the studied ori-*GOI*) is lower than the predicted value of (1 − *e*^−λ^) × 100 = 18%, with *λ* = 0.2 at 10 pM DNA, if one assumes that all liposomes with at least one DNA copy would give a signal. This discrepancy suggests that (i) some DNA molecules are transcriptionally inactive (Nourian and Danelon 2013), thus lowering the apparent *λ*, (ii) only liposomes with particularly high concentration of amplified DNA or reporter protein cross the fluorescence detection threshold in our flow cytometer-based activity assay, (iii) the encapsulation of input DNA does not follow a Poisson distribution, or (iv) the heterogeneous size distribution of the liposomes (approximately lognormal) (Blanken et al., 2019) shifts *λ* to lower values than expected if one assumes a constant size of 4 μm diameter. We suspect that under some conditions (Fig. 2d and Fig. 5h with ori-*yfp* and ori-*pssA*, respectively) competition for resources during *p2* and *p3* expression may limit the yield of synthesized POI in expCADGE. Therefore, optimization of the concentration of the *p2-p3* expression plasmid might be necessary for effective channeling of resources toward expression of POI. This drawback may be alleviated to some extent by using purified DNAP and TP (purCADGE, see Fig 2 and Fig 5). Another limitation is the current lack of commercial availability of some of the required components, such as purified TP, SSB, or DSB. However, TP (together with DNAP) can be expressed from a plasmid in situ (expCADGE), while Φ29 SSB might in principle be replaced with a commercially available alternative (such as *E. coli* SSB), provided that it is shown to be compatible. Overall, we recommend that the optimality of expCADGE vs purCADGE, as well as the optimal DSB and *p2-p3* expression plasmid concentrations should be determined on a case-by-case basis for each GOI. We also believe that the yield of POI production per DNA template could be further improved through buffer optimization, in particular the concentrations of magnesium and potassium glutamate (Borkowski et al., 2020), tRNAs and NTPs (Sakatani et al., 2015).

PURE*frex2*.*0* in vitro transcription-translation system was used here in its standard composition. However, other promoters than T7, such as the bacteriophage SP6 (Ishikawa et al., 2004; Blanken et al., 2020), T3 (Kobori et al., 2013) or native *E. coli* (Maddalena et al., 2016) promoters, could also be used in combination with their cognate RNA polymerase supplied in the reaction mixture. One challenge may reside in the management of collision events between the Φ29 DNA polymerase and the RNA polymerases originating from different organisms (Elías-Arnanz and Salas, 1997, 1999)(Elías-Arnanz and Salas 1997 and 1999). Moreover, cell lysates, especially from *E. coli* (Marshall and Noireaux, 2018), could be utilized as a cheaper cell-free expression system, in particular for biomanufacturing purposes (Silverman et al., 2020). The extract could be modified to avoid the degradation of linear PCR fragments by exonucleases, for instance by supplementing inhibitors of RecBCD (ExoV), such as GamS protein (Sitaraman et al., 2004)or *χ*-DNA oligonucleotides (Marshall et al., 2017) or using Δ*RecBCD E. coli* strains (Batista et al., 2022). Alternatively, purified TP-bound DNA (van Nies et al., 2018) could be used as a template in cell lysates, assuming that the parental TP hinders exonuclease digestion. Application of CADGE in eukaryotic cell extracts – e.g. from insect cells, wheat germ, rabbit reticulocytes, and human cells – might be useful for the production and engineering of proteins with post-translational modifications, such as glycosylation and phosphorylation. While protein yields may remain low even with clonal gene amplification compared to *E. coli*-based cell-free systems, the increased amount of DNA may be sufficient for the recovery of interesting gene variants. In general, codon usage of GOI may be optimized for the chosen cell-free translation system, which should not influence much the DNA replication efficacy given the high template tolerance of the Φ29 DNAP.

In the shown examples, the genotype-to-phenotype coupling was established using phospholipid vesicles. Liposomes are uniquely suited for cell-free evolution of peripheral and transmembrane proteins (Fujii et al., 2013; Uyeda et al., 2016), and for their tunable membrane permeability, which could be relevant to assay the activity of POI through external addition of substrates or cofactors. The lipid-coated bead approach for liposome production (Nourian et al., 2012) was chosen for its simplicity as it does not require specialized equipment, for the easy storage and distribution across laboratories of pre-assembled lipid films deposited on glass microbeads, and for its high biocompatibility due to the absence of organic solvent. Other liposome preparation methods, such as enhanced continuous droplet interface crossing encapsulation (Van de Cauter et al., 2021) and double-emulsion microfluidics (Gonzales et al., 2022), could in principle be utilized as well.

Other types of microcompartments can also potentially be combined with CADGE: water-in-oil emulsion droplets (Fallah-Araghi et al., 2012), microfabricated chambers (Zhang et al., 2019), and peptide-based compartments (Vogele et al., 2018). Emulsion droplets are particularly appealing for their high monodispersity and because they have already been integrated in microfluidic-based screening platforms for directed evolution of water-soluble enzymes (Obexer et al., 2017; Holstein et al., 2021).

Application of gene expressing liposomes empowered with clonal amplification is also relevant to build a synthetic cell from the ground-up. When applied to essential genes, CADGE-assisted directed evolution might accelerate the optimization of individual cellular modules and their integration to achieve higher level functions (Abil and Danelon, 2020). Considering the excellent processivity of Φ29 DNAP (Blanco et al., 1989; Rodríguez et al., 2005), the application of CADGE to long synthetic genomes can reasonably be envisaged. Through the example of TP (Fig. 3), we showed the implementation of a positive feedback loop, where the GOI can itself assist in its own amplification, thereby circumventing the need for screening. This reaction scheme may in principle be expanded to self-amplification of polymerases and gene circuits based on DNA polymerization, such as in compartmentalized self-replication (Ghadessy et al., 2001) and compartmentalized partnered replication (Ellefson et al., 2014). Moreover, self-organization and catalytic activity of the peripheral membrane proteins MinD and PssA were detectable by isothermal DNA amplification from clonal amounts. This strategy might be particularly useful for the in vitro evolution of cellular functions starting from a single copy of ori-*GOI* library variants encapsulated in liposomes. The replicating template may contain single or multiple genes encoding entire pathways and multiprotein complexes. For instance, application of CADGE to phospholipid-synthesizing enzymes of the Kennedy pathway located upstream (PlsB, PlsC and CdsA) and downstream (Psd) of PssA could aid in optimizing synthetic cell growth through directed evolution.

Finally, we anticipate that performing CADGE under mutagenic conditions could extend its utility for in situ library production. For example, a mutator DNA polymerase (de Vega et al., 1996) or mutagenic factors, such as Mn^2+^ and dNTP analogues, could be employed for genetic diversification directly within liposomes, bypassing the step of external gene library preparation. Such an error-prone CADGE strategy might be particularly interesting for introducing random mutations across long (> 10 kbp) DNA templates, for instance large synthetic genomes for the evolutionary construction of a minimal cell (Abil and Danelon, 2020).

## Methods

### Buffers and solutions

All buffers and solutions were made using Milli-Q grade water with 18.2 MΩ resistivity (Millipore, USA). Chemicals were purchased from Sigma-Aldrich unless otherwise indicated.

### Construct design

G365 (pUC-ori-YFP) was constructed by subcloning of the YFP gene (amplified by primers 1106 ChD/1107 ChD from plasmid G79) into Φ29 origins-containing vector G96 (van Nies et al., 2018) (amplified by primers 1104 ChD /1105 ChD) via the Gibson Assembly method (Gibson et al., 2009). G368 (pUC-ori-pssA) was cloned by subcloning of the *pssA* gene (amplified by primers 1115 ChD /1116 ChD from plasmid G149) into Φ29 origins-containing vector G96 (amplified by primers 1104 ChD /1105 ChD) via the Gibson Assembly method. Plasmid G338 (pUC-ori-TP) was obtained as a result of subcloning the fragment ori-p2p3, which was PCR-amplified from plasmid G95 (plasmid encoding for ori-*p2-p3*, (van Nies et al., 2018)) using the primers 961 ChD /962 ChD (with overhangs containing KpnI and HindIII restriction sites) into the KpnI-HindIII-linearized pUC19 vector, during which a spontaneous recombination event flipped out the entire (t7)promoter-p2-(vsv)terminator fragment, only leaving the shorter oriL-(t7)promoter-p3-(t7)terminator-oriR insert. G437 (pUC-ori-minD) was obtained by subcloning the *minD* gene (amplified by primers 91 ChD /397 ChD from plasmid pUC57-MinD) (Godino et al., 2019) into the Φ29 origins-containing vector G365 (amplified by primers 535 ChD/562 ChD) via the Gibson Assembly method. The cloning of G85 (pUC57-DNAP) was previously reported (van Nies et al., 2018). All the plasmids were cloned by heat-shock transformation of *E. coli* Top10 strain, and plasmids were extracted from individual cultures outgrown in LB/ampicillin (50 μg/mL) using the PURE Yield Plasmid Miniprep kit (Promega). Individual clones were screened and confirmed by Sanger sequencing at Macrogen-Europe B.V. Primer sequences and plasmid descriptions can be found in the Supplementary Table 1 and Table 2.

Linear DNA templates were prepared by PCR using 5’-phosphorylated primers (491 ChD /492 ChD). Reactions were set up in 100 μL volume, 500 nM each primer, 200 μM dNTPs, 10 pg/μL DNA template, and 2 units of Phusion High-Fidelity DNA Polymerase (NEB) in HF Phusion buffer, and thermal cycling was performed as follows: 98 °C 30 sec initial denaturation, and thermal cycling at (98 °C for 5 sec, 72 °C for 90 sec) × 20, and final extension at 72 °C for 5 min. Extra care was taken to not over-amplify the DNA by too many thermal cycles, as it was found to adversely affect the quality of purified DNA. The amplified PCR fragments were purified using QIAquick PCR purification buffers (Qiagen) and RNeasy MinElute Cleanup columns (Qiagen) using the manufacturer’s guidelines for QIAquick PCR purification, except for longer pre-elution column drying step (4 min at 10,000 g with open columns), and elution with 14 μL ultrapure water (Merck Milli-Q) in the final step. The purified DNA was quantified by Nanodrop 2000c spectrophotometer (Isogen Life Science) and further analyzed for size and purity by gel electrophoresis.

### Purification of DNAP, TP, SSB, DSB, LactC2-eGFP and LactC2-mCherry

Purified Φ29 DNA replication proteins were produced as described in (van Nies et al., 2018). Stock concentrations and storage buffers are: DNAP (320 ng/μL in 50 mM Tris, pH 7.5, 0.5 M NaCl, 1 mM EDTA, 7 mM β-mercaptoethanol (BME), 50% glycerol), TP (400 ng/μL in 25 mM Tris, pH 7.5, 0.5M NaCl, 1 mM EDTA, 7 mM BME, 0.025% Tween 20, 50% glycerol), SSB (10 mg/mL in 50mM Tris, pH 7.5, 60 mM ammonium sulfate, 1 mM EDTA, 7 mM BME, 50% glycerol), DSB (10 mg/mL in 50 mM Tris, pH 7.5, 0.1 M ammonium sulfate, 1 mM EDTA, 7 mM BME, 50% glycerol). The proteins were aliquoted and stored at −80 °C. The DNAP and TP proteins were diluted before immediate use into PURE*frex2*.*0* solution I (GeneFrontier). Both LactC2-eGFP and LactC2-mCherry genes were cloned into pET11 vector, under control of the T7-LacO promotor and in frame with an N-terminal His-tag. LactC2-mCherry was expressed in *E. coli* BL21(DE3) (NEB) and LactC2-eGFP protein was expressed in *E. coli* strain ER2566 (NEB). Overnight pre-cultures were prepared in Luria Broth (LB) medium containing 50 μg/mL ampicillin. The overnight cultures were diluted 1:100 in fresh LB medium with 50 μg/mL ampicillin and incubated at 37 °C while shaking, until an OD_600_ of 0.4-0.6 was reached. Protein expression was induced by adding 1 mM isopropyl β-D-1-thiogalactopyranoside. The cells were incubated at 26 °C for 4 h or overnight at 16 °C while shaking, and harvested at 4,000 × g for 15 min. Pellet of 1 L cells was resuspended in 10 mL lysis buffer (50 mM HEPES-KOH, pH 7.5, 500 mM NaCl, 10% glycerol). The cells were disrupted by sonication on ice, using 7 pulses of 30 seconds and 1 min intervals, with an amplitude of 40%. The cell suspension was centrifuged for 30 min at 30,000 × g at 4 °C to remove the cell debris. To the cell-free extract, 10 mM imidazole and SetIII protease inhibitor-EDTA-free (1:1000 dilution, Calbiochem) were added. The proteins were purified with HisPure Ni-NTA resin (ThermoScientific). The Ni-NTA (∼3 mL) was equilibrated with buffer (50 mM HEPES-KOH, 500 mM NaCl, 10% glycerol, 10 mM imidazole, pH 7.5). The cell-free extract was mixed with the equilibrated resin and incubated for 1 to 16 h while tumbling in the cold room. After incubation the resin with bound protein was transferred into a gravity column, the unbound fraction was removed by gravity and subsequently the resin was washed with 20 equivalent volume wash buffer (50 mM HEPES-KOH, 500 mM NaCl, 10% glycerol, 40 mM imidazole, pH 7.5). The protein was eluted with 5 mL elution buffer (50 mM HEPES-KOH, 500 mM NaCl, 500 mM imidazole, 10% glycerol, pH 7.5) and fractions of ∼1 mL were collected. The fluorescent fractions were pooled together and buffer exchanged with storage buffer (50 mM HEPES-KOH, pH 7.5, 150 mM NaCl, 10% glycerol) using a 10-MWCO Amicon Ultra-15 centrifugal filter unit (Merck). The concentration of the protein was determined with a Bradford assay.

### CADGE in bulk reactions

Bulk reactions were set up in PURE*frex2*.*0* (GeneFrontier). A 20-μL reaction consisted of 10 μL solution I, 1 μL solution II, 2 μL solution III, 20 mM ammonium sulfate, 300 μM dNTPs, 375 μg/mL purified Φ29 SSB protein, 105 μg/mL purified Φ29 DSB protein, 0.6 units/μL of Superase·In RNase inhibitor (Ambion), 10 pM target DNA and either plasmid DNA (2 nM plasmid G85 encoding for the *p2* gene in ori-*p3* clonal amplification experiments or 1 nM G340 encoding for *p2* and *p3* genes in ori-*yfp*, ori-*minD*, and ori-*pssA* experiments) or 3 ng/μL each purified Φ29 DNAP and TP. Reactions were incubated in a nuclease-free PCR tube (VWR) in a ThermalCycler (C1000 Touch, Biorad) at a default temperature of 30 °C. Incubation time was indicated when appropriate.

To analyse the reactions by gel electrophoresis, 10-μL reaction was treated with 0.2 mg/mL RNase A (Promega), 0.25 units RNase One (Promega) at 30 °C for 2 h, followed by 1 mg/mL Proteinase K (Thermo Scientific) at 37 °C for 1 h, and column-purified using the QIAquick PCR purification buffers (Qiagen) and RNeasy MinElute Cleanup columns (Qiagen) using the manufacturer’s guidelines for QIAquick PCR purification, except for longer pre-elution column drying step (4 min at 10,000 g with open columns), and elution with 14 μL ultrapure water (Merck Milli-Q) in the final step. A fraction (6 μL) of the eluate was mixed with an equal volume of 6× purple gel loading dye (NEB) and loaded in 1% agarose gel with ethidium bromide, following which DNA was separated using an electrophoresis system (Bio-Rad). The BenchTop 1-kb DNA Ladder (Promega) was used to estimate the size of DNA.

### Mass spectrometry

LC-MS/MS analysis with QconCATs was employed for the absolute quantification of de novo synthesized proteins in bulk PURE reactions. Pre-ran PURE reaction solutions were mixed with one third volume of heavily labeled QconCAT(^15^N) (Doerr et al., 2021) in a 50 mM Tris (pH 8.0) buffer containing 1 mM CaCl_2_. The samples were then incubated at 90 °C for 10 min and cooled down to 4 °C. Trypsin was then added at a 250 μg/mL final concentration and digestion incubation was carried out overnight at 37 °C. The trypsin digested samples were treated with TFA 10% and centrifuged for 10 min. The supernatant was then transferred to a glass vial with a small insert for LC-MS/MS analysis. Measurements were performed on a 6460 Triple Quad LCMS system (Agilent Technologies, USA) using Skyline software (MacLean et al., 2010). Samples of 5.5 μL were injected per run into an ACQUITY UPLC Peptide CSH C18 Column (Waters Corporation, USA). The peptides were separated in a gradient of buffer A (25 mM formic acid in Milli-Q water) and buffer B (50 mM formic acid in acetonitrile) at a flow rate of 500 μL per minute and at a column temperature of 40 °C. The column was initially equilibrated with 98% buffer A. After sample injection, buffer A gradient was changed to 70% (over the first 20 min), 60% (over the next 4 min), and 20% (over the next 30 sec). This final ratio was conserved for another 30 sec and the column was finally flushed with 98% buffer A to equilibrate it for the next run. The selected peptides and their transitions for both synthesized proteins and heavily labeled QconCATs were measured by multiple reaction monitoring (MRM). The recorded LC-MS/MS data was analyzed with Skyline for fraction calculation between unlabeled and labeled peptides (^14^N/^15^N ratio) on both cell-free core/produced proteins and the initially added QconCATs. With these fraction values, and considering the regular concentration of core ribosomal peptides within PURE system (2 μM), we could estimate the concentration of the cell-free expressed proteins using the following equation:

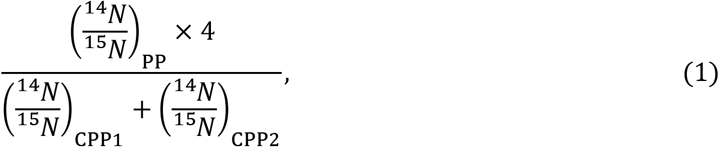

where PP refers to the detected peptide of produced protein, CPP1 refers to the detected peptide 1 of the core ribosomal protein (GVVVAIDK), and CPP2 refers to the detected peptide 2 of the core ribosomal protein (VVGQLGQVLGPR). MS/MS measurement details for each of the analyzed proteins can be found in Supplementary Table 3.

### In-vesiculo protein expression

The procedure was adapted from (van Nies et al., 2018) with minor modifications. To prepare lipid-coated beads as precursors of liposomes with the standard lipid composition, in a 5-mL round-bottom glass flask, a primary lipid mixture was prepared consisting of DOPC (50.8 mol%), DOPE (35.6 mol%), DOPG (11.5 mol%), and 18:1 cardiolipin (2.1 mol%). The resulting mixture was additionally spiked with DSPE-PEG(2000)-biotin (1 mass%) and DHPE-Texas Red (0.5 mass%) for a total mass of 2 mg. Finally, the lipid mixture was complemented with 25.4 μmol of rhamnose (Sigma-Aldrich) dissolved in methanol. To prepare liposomes containing CDP-DAG, the primary lipid mixture composition was modified as following: DOPC (47.5 mol%), DOPE (34.2 mol%), DOPG (11.4 mol%), 18:1 cardiolipin (1.9 mol%), and CDP-DAG (5 mol%), with additional DSPE-PEG(2000)-biotin (1 mass%) and, if indicated, DHPE-TexasRed (0.5 mass%) for a total mass of 2 mg. All lipids were purchased at Avanti Polar Lipids and dissolved in chloroform, except the DHPE-Texas Red (Invitrogen). Finally, 600 mg of 212–300-μm glass beads (Sigma-Aldrich) were added to the lipid/rhamnose solution, and the organic solvent was removed by rotary evaporation at 200 mbar for 2 h at room temperature (rotary evaporator, Heidolph), followed by overnight desiccation. The dried lipid-coated beads were stored under argon at −20 °C until use. A 20-μL PURE*frex2*.*0* reaction solution was assembled from 10 μL buffer solution, 1 μL enzyme solution, 2 μL ribosome solution, indicated amount of input DNA template in RNase-free Milli-Q water. To the well-mixed reaction, 10 mg lipid-coated beads, additionally pre-dessicated for at least 30 min before use, were added. The 1.5-mL Eppendorf tube containing the bead-PURE*frex2*.*0* mixture was gently rotated on an automatic tube rotator (VWR) at 4 °C along its axis for 30 min for uniform liposome swelling. The mixtures were then subjected to four freeze/thaw cycles (5 sec in liquid nitrogen followed by 10 min on ice). From this step onwards, the liposome suspension was handled gently and only with cut pipette tips to prevent liposome breakage. Finally, 10 μL of the supernatant liposome suspension (the beads sediment to the bottom of the tube) was transferred to a PCR tube, where it was mixed with 0.5 units of DNase I (NEB). The reactions were incubated at 30 °C in a thermocycler for the indicated time periods.

### In-vesiculo clonal amplification and expression of genes

The liposome suspensions were assembled as above, except that the necessary CADGE components were pre-mixed with the PURE*frex2*.*0* solution prior to the addition of the lipid-coated beads and swelling. The following compounds were supplemented (all final concentrations): 20 mM of ammonium sulfate, 0.75 U/μL SUPERase (Ambion), 10-50 pM template DNA (as indicated), 375 μg/μL purified SSB, 21-52.5 μg/μL of purified DSB, 1-3 ng/μL each of purified Φ29 DNAP and TP proteins or 250 pM of the *p2-p3* encoding G340 plasmid, and 300 μM of PCR Nucleotide mix (Promega). The liposome suspensions were incubated at 30 °C in a thermocycler for the indicated time periods.

### Quantitative PCR

Upon completion of the bulk or in-liposome CADGE reactions at 30 °C, 2-μL samples were harvested, heated at 75 °C in the thermocycler for 15 min to inactivate the DNase I, and diluted 100-fold in Milli-Q water prior to addition to the qPCR mixtures. Ten microliter reactions consisted of PowerUP SYBR Green Master Mix (Applied Biosystems), 400 nM each primer (1121 ChD/1122 ChD for *yfp*, 980 ChD/981 ChD for *p3*, 1125 ChD/1126 ChD for *pssA*, 1208 ChD/1209 ChD for *minD*), and 1 μL of diluted sample. The thermal cycling and data collection were performed on Quantstudio 5 Real-Time PCR instrument (Thermo Fisher), using the thermal cycling protocol 2 min at 50 °C, 5 min at 94 °C, (15 sec at 94 °C, 15 sec at 56 °C, 30 sec at 68 °C) × 45, 5 min at 68 °C, followed by melting curve from 65 °C to 95 °C. The concentration of nucleic acids was calibrated using 10-fold serial dilutions of corresponding standard DNA templates ranging from 1 fM to 1 nM. Data were analysed using the Quantstudio Design and Analysis software v1.4.3 Software (Thermo Fisher).

### Flow cytometry

The liposome suspension (1-3 μL) was diluted in 300 μL (final volume) PB buffer consisting of 20 mM HEPES-KOH, pH 7.6, 180 mM potassium glutamate, and 14 mM magnesium acetate. To remove any remaining beads or large debris, the diluted liposome suspension was gently filtered through the 35-μm nylon mesh of the cell-strainer cap from the 5-mL round-bottom polystyrene test tubes (Falcon). When indicated, DsGreen (Lumiprobe) dye was added at a 1:100,000 stock concentration to stain dsDNA, or Acridine Orange (6 μM) and LactC2-mCherry protein (300 nM) were added to stain the liposome membrane and phosphatydilserine, respectively. The mixture was incubated for 1 h at room temperature to equilibrate binding. Liposomes were screened with the FACSCelesta flow cytometer (BD Biosciences) using the 488-nm laser and 530/30 filter for detection of DsGreen, GFP, YFP, or Acridine Orange, and the 561-nm laser and 610/20 filter for detection of PE-Texas Red or mCherry. The following acquisition parameters were used: photon multiplier tube voltages set at 375 V for forward scatter, 260 V for side scatter (SSC), DsGreen detection at 500 V, PE-Texas Red detection at 300-370 V, YFP detection at 550 V, GFP detection at 700 V, mCherry detection at 370 V, Acridine Orange detection at 400 V, threshold for SSC at 200 V, sample flow 1 (∼1000 events/s), injection volume 50-200 μL, recording of 10-100,000 total events. Data were analyzed using Cytobank (https://community.cytobank.org/). Raw data was pre-processed as described in Fig. S8 to filter out possible aggregates and debris.

### Confocal microscopy and image analysis

A custom-made glass imaging chamber was functionalized with BSA-biotin:BSA and Neutravidin as previously described (Blanken et al., 2019). The liposome suspension (3-7 μL) was supplemented with PB buffer to a maximum volume of 7 μL and transferred into the functionalized chamber. The LactC2-GFP probe was used at a final concentration of ∼260 nM.

After 30 to 60 min incubation at room temperature to let the liposomes sediment, the sample was imaged with a Nikon A1R Laser scanning confocal microscope equipped with a ×100 objective and operated via the NIS Elements software (Nikon). The laser lines 488 nm (for MinC-eGFP), 514 nm (for YFP) and 561 nm (for DHPE-Texas Red and LactC2-mCherry) were used in combination with appropriate emission filters. The position of the focal plane was manually adjusted to image as many liposomes as possible across their equatorial plane. Image analysis was performed using ImageJ (https://imagej.nih.gov/ij/) and an in-house developed code, called SMELDit, which enables the identification of individual liposomes, as well as the quantification of fluorescence signals at the membrane and in the lumen.

### Mock enrichment of *p3* gene

The linear DNA constructs ori-*p3* and ori-*p6* were mixed at a 1:1 molar ratio for a total DNA concentration of either 10 pM or 50 pM in PURE*frex2*.*0* solutions containing 20 mM ammonium sulfate, 300 μM dNTPs, 375 μg/mL purified SSB, 105 μg/mL purified DSB, and 0.6 units/μL of Superase·In RNase inhibitor. The reactions were also supplemented with 2 nM of plasmid DNA encoding for Φ29 DNAP (G85 plasmid). The well-mixed solution was encapsulated in liposomes as described above. Then, 5 μL of bead-free liposome suspension was transferred to a PCR tube, where it was mixed with 0.25 units of DNase 1 (Thermo Scientific), and incubated at 30 °C for 16 h. Upon completion, 2-μL samples were harvested from both + and – dNTPs reactions for quantitative PCR as described above.

### Mock enrichment of *yfp* gene

The linear DNA constructs ori-*yfp* and ori-*minD* were mixed at 1:10 molar ratio (1 pM ori-*yfp* and 9 pM ori-*minD* final concentrations) in either gene expression solution (PURE*frex2*.*0*: 50% v/v solution I, 5% v/v solution II, and 10% v/v solution III supplemented with 0.6 units/μL of Superase·In RNase inhibitor) or gene expression-coupled replication solution (PURE*frex2*.*0* with an addition of 20 mM ammonium sulfate, 300 μM dNTPs, 375 μg/mL purified SSB, 52.5 μg/mL purified DSB, 3 ng/μL purified Φ29 DNAP, 3 ng/μL purified TP, and 0.6 units/μL of Superase·In RNase inhibitor). The well-mixed solution was encapsulated in liposomes as described above. Then, 10 μL of bead-free liposome suspension was transferred to a PCR tube, where it was mixed with 0.5 units of Proteinase K (Thermo Sceintific), and incubated at 30 °C for 16 h. Three microliter of liposome suspension was mixed with 497 μL PB buffer and filtered through the 35-μm nylon mesh of the cell-strainer cap from the 5-mL round-bottom polystyrene test tubes (Falcon).

Fluorescence-activated cell sorting was conducted on FACSMelody (BD Biosciences). Lasers PE-CF594(YG) and FITC-BB515, 100-μm nozzle, 23.14 PSI pressure and 34.2 kHz drop frequency were used. Photon multiplier tube voltages applied were 320 V for forward scatter, 455 for side scatter, 337 V for Texas Red, and 673 V for GFP, and a threshold of 359 V at the side scatter was applied. Liposomes with 1% highest YFP signal were sorted out from liposomes prepared in gene expression solution (“all-gate”), and the same gate was applied to the liposomes prepared in gene expression-coupled replication solution or an adjusted gate including only 0.2% highest YFP signal (“high-gate”). Around 50,000 (low-gate) or 10,000 (high-gate) liposomes were sorted into a 1.5 mL Eppendorf tube. Liposomes from the “all-gate” were further concentrated by centrifugation at 12,000 g for 3 min, and removing three fourth of the supernatant volume. The proteinase K was heat inactivated at 95 °C for 5 min.

The DNA contained in sorted liposomes was used as a template for PCR amplification using phosphorylated primers (ChD 491/ChD 492). Reactions were set up in 100 μL volume, 300 nM each primer, 400 μM dNTPs, 10 μL sorted, heat-inactivated liposome solution, and 2 units of KOD Xtreme Hotstart DNA polymerase in Xtreme buffer, and thermal cycling was performed as follows: 2 min at 94 °C for polymerase activation, and thermal cycling at (98 °C for 10 sec, 65 °C for 20 sec, 68 °C for 1.5 min) × 30. The amplified PCR fragments were purified using QIAquick PCR purification buffers (Qiagen) and RNeasy MinElute Cleanup columns (Qiagen) using the manufacturer’s guidelines for QIAquick PCR purification, except for longer pre-elution column drying step (4 min at 10,000 g with open columns), and elution with 14 μL ultrapure water (Merck Milli-Q) in the final step. The purified DNA was quantified by the Nanodrop 2000c spectrophotometer (Isogen Life Science).

### Statistical analysis of DNA occupancy

The probability that a liposome contains *k* molecules of DNA (*k* = 0, 1, 2, 3, …) according to a Poisson distribution is:

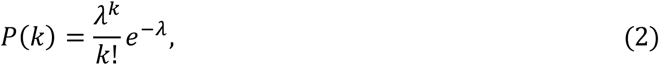

where *λ* is the expected average number of input DNA molecules per liposome. It can be calculated as a function of the diameter *d* of the liposomes and the bulk concentration *C* of input DNA templates, as:

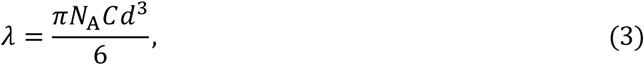

where *N*_*A*_ is the Avogadro constant. A CADGE reaction may occur in a liposome if one or more copies of linear DNA template is encapsulated, whose corresponding probability is given by:

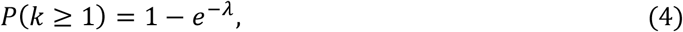

With expCADGE, the concentration of *p2-p3*-plasmid largely exceeds that of ori-*GOI*, such that only the concentration of ori-*GOI* template limits the percentage of liposomes exhibiting CADGE:

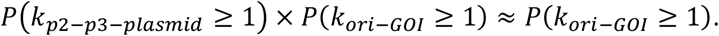

## Data and code availability statement

Flow cytometry data were analyzed using Cytobank (https://community.cytobank.org/). MATLAB scripts and a user manual for SMELDit are made available upon request.

## Acknowledgments

We are grateful to Margarita Salas and Mario Mencía Caballero (Universidad Autónoma de Madrid), and Alicia del Prado and Miguel de Vega (Centro de Biología Molecular Severo Ochoa, Madrid) for kindly providing us with the purified DNAP, TP, SSB, and DSB proteins. We thank Yannick Rondelez and Thibault Di Meo (ESPCI, Paris) for fruitful discussions. We also thank Ilja Westerlaken for the purification of LactC2-mCherry and LactC2-eGFP proteins, Elisa Godino for helping with the preparation and imaging of MinD-containing liposome samples. We are grateful to Duco Blanken, Flora Yang and Anne Doerr for their assistance with the mass spectrometry experimental set up. Finally, we are thankful to Tess Bevers for assistance with cloning of the ori-*p3* plasmid and Mats van Tongeren for developing SMELDit. This project has received funding from the European Union’s Horizon 2020 research and innovation programme under the Marie Skłodowska-Curie grant agreement No 707404, and from the Netherlands Organization for Scientific Research (NWO/OCW) via the “BaSyC – Building a Synthetic Cell” Gravitation grant (024.003.019).

