## Supplementary Information for "Clonal amplification-enhanced gene expression for cell-free directed evolution"

**Supplementary Table 1.** DNA primer sequences and purpose.

| Name | Purpose | Sequence | Modifications |
|---|---|---|---|
| 1106 ChD | YFP gene subcloning | CCGTTTAGAGGCCCCAAGGG | none |
| 1107 ChD | YFP gene subcloning | CTTCGTCTGTGTCGCATGTGAaATTAATACGACTCACTATAGGGAGACCACAACG | none |
| 1104 ChD | amplification of vector | TAGCATAACCCCTTGGGGC | none |
| 1105 ChD | amplification of vector | CCTATAGTGAGTCGTATTAATtTCACATGCGAC | none |
| 1115 ChD | pssA gene subcloning | CTTCGTCTGTGTCGCATGTGAaATTAATACGACTCACTATAGGGGAATTGTGAGC | none |
| 1116 ChD | pssA gene subcloning | AACCCCTCAAGACCCGTTTAGAG | none |
| 961 ChD | TP gene cloning | acgtggtaccAAAGTAAGCCCCCACCCTCACATG | none |
| 962 ChD | TP gene cloning | agctaagcttAAAGTAGGGTACAGCGACAACATACAC | none |
| 491 ChD | preparative PCR for IVTTR | 5-phos/AAAGTAAGCCCCCACCCTCACATG | 5'-phosphorylation |
| 492 ChD | preparative PCR for IVTTR | 5-phos/AAAGTAGGGTACAGCGACAACATACAC | 5'-phosphorylation |
| 1121 ChD | YFP detection | TGCAACTGGCTGACCACTAC | none |
| 1122 ChD | YFP detection | AATGATTGTCCGGCAGCAGA | none |
| 980 ChD | p3 detection | ACGGCTGAAATTGACATCCCG | none |
| 981 ChD | p3 detection | CCAGGCGTTGAACTTCTTTGG | none |
| 1125 ChD | pssA-qPCR-F | AACAGGATGACGGTGGCAAA | none |
| 1126 ChD | pssA-qPCR-R | GGAACATCTACGCCCGGATT | none |
| 1208 ChD | MinD detection | CGCGACTCTGACCGTATTT | none |
| 1209 ChD | MinD detection | AGCATGTCACCTCTGCTTAC | none |

**Supplementary Table 2.** Plasmid DNA description.

| plasmid name | plasmid description |
|---|---|
| **G365** | Contains the DNA unit for the expression of YFP fluorescence protein. Transcription is regulated by a T7 promoter and T7 terminator sequences. The entire CDS unit is placed in between right and left origins of replication from the Φ29 DNA replication machinery. |
| **G368** | Contains the DNA unit for the expression of the phospholipid biosynthesis protein PssA. Transcription is regulated by a T7 promoter and T7 terminator sequences. The entire CDS unit is placed in between right and left origins of replication from the Φ29 DNA replication machinery. |
| **G437** | Contains the DNA unit for the expression of MinD protein. Transcription is regulated by a T7 promoter and T7 terminator sequences. The entire CDS unit is placed in between right and left origins of replication from the Φ29 DNA replication machinery. |
| **G338** | Contains the DNA unit for the expression of the Φ29 terminal protein TP. Transcription is regulated by a T7 promoter and T7 terminator sequences. The entire CDS unit is placed in between right and left origins of replication from the Φ29 DNA replication machinery. |
| **G85** | Contains the DNA unit for the expression of Φ29 DNA polymerase. Transcription is regulated by a T7 promoter and vsv terminator sequences. |
| **G95** | Contains the DNA sequence for the expression of DNAP and TP. Each protein expression is independently regulated by a T7 promoter and a terminator sequence. DNAP unit uses a vsv terminator. TP unit utilizes a T7 terminator sequence. The entire CDS encoding for DNAP and TP is placed in between right and left origins of replication from the Φ29 DNA replication machinery. |

**Supplementary Table 3.** Transitions of the MS/MS measurements for the proteolytic peptides of the indicated proteins.

| Protein | Compound name | Precursor ion *(m/z)* | Product ion *(m/z)* | Collision energy (eV) | Accelerator voltage (eV) | Ion name |
|---|---|---|---|---|---|---|
| PSSA | DLQSIADYPVK.light | 624.8272 | 805.4454 | 20.4 | 4 | y7 |
| PSSA | DLQSIADYPVK.light | 624.8272 | 692.3614 | 20.4 | 4 | y6 |
| PSSA | DLQSIADYPVK.light | 624.8272 | 506.2973 | 20.4 | 4 | y4 |
| PSSA | DLQSIADYPVK.light | 624.8272 | 343.2340 | 20.4 | 4 | y3 |
| PSSA | DLQSIADYPVK.light | 624.8272 | 357.1769 | 20.4 | 4 | b3 |
| PSSA. QconCAT | DLQSIADYPVK.heavy | 631.3079 | 813.4217 | 20.4 | 4 | y7 |
| PSSA. QconCAT | DLQSIADYPVK.heavy | 631.3079 | 699.3406 | 20.4 | 4 | y6 |
| PSSA. QconCAT | DLQSIADYPVK.heavy | 631.3079 | 511.2825 | 20.4 | 4 | y4 |
| PSSA. QconCAT | DLQSIADYPVK.heavy | 631.3079 | 347.2221 | 20.4 | 4 | y3 |
| PSSA. QconCAT | DLQSIADYPVK.heavy | 631.3079 | 361.1650 | 20.4 | 4 | b3 |
| YFP | FEGDTLVNR.light | 525.7644 | 903.4530 | 17.3 | 4 | y8 |
| YFP | FEGDTLVNR.light | 525.7644 | 774.4104 | 17.3 | 4 | y7 |
| YFP | FEGDTLVNR.light | 525.7644 | 717.3890 | 17.3 | 4 | y6 |
| YFP | FEGDTLVNR.light | 525.7644 | 602.3620 | 17.3 | 4 | y5 |
| YFP | FEGDTLVNR.light | 525.7644 | 501.3144 | 17.3 | 4 | y4 |
| YFP | FEGDTLVNR.light | 525.7644 | 449.1667 | 17.3 | 4 | b4 |
| YFP. QconCAT | FEGDTLVNR.heavy | 532.2451 | 915.4175 | 17.3 | 4 | y8 |
| YFP. QconCAT | FEGDTLVNR.heavy | 532.2451 | 785.3778 | 17.3 | 4 | y7 |
| YFP. QconCAT | FEGDTLVNR.heavy | 532.2451 | 727.3593 | 17.3 | 4 | y6 |
| YFP. QconCAT | FEGDTLVNR.heavy | 532.2451 | 611.3354 | 17.3 | 4 | y5 |
| YFP. QconCAT | FEGDTLVNR.heavy | 532.2451 | 509.2906 | 17.3 | 4 | y4 |
| YFP. QconCAT | FEGDTLVNR.heavy | 532.2451 | 453.1548 | 17.3 | 4 | b4 |
| Ribosomal protein S4 (YFP quantification) | LSDYGVQLR.light | 525.7826 | 850.4417 | 17.3 | 4 | y7 |
| Ribosomal protein S4 (YFP quantification) | LSDYGVQLR.light | 525.7826 | 735.4148 | 17.3 | 4 | y6 |
| Ribosomal protein S4 (YFP quantification) | LSDYGVQLR.light | 525.7826 | 572.3515 | 17.3 | 4 | y5 |
| Ribosomal protein S4 (YFP quantification) | LSDYGVQLR.light | 525.7826 | 635.3035 | 17.3 | 4 | b6 |
| Ribosomal protein S4. QconCAT (YFP quantification) | LSDYGVQLR.heavy | 525.7826 | 861.4091 | 17.3 | 4 | y7 |
| Ribosomal protein S4. QconCAT (YFP quantification) | LSDYGVQLR.heavy | 525.7826 | 745.3851 | 17.3 | 4 | y6 |
| Ribosomal protein S4. QconCAT (YFP quantification) | LSDYGVQLR.heavy | 525.7826 | 581.3248 | 17.3 | 4 | y5 |
| Ribosomal protein S4. QconCAT (YFP quantification) | LSDYGVQLR.heavy | 525.7826 | 641.2857 | 17.3 | 4 | b6 |
| Ribosomal protein L6 (YFP quantification) | APVVVPAGVDVK.light | 575.8452 | 883.5247 | 18.9 | 4 | y9 |
| Ribosomal protein L6 (YFP quantification) | APVVVPAGVDVK.light | 575.8452 | 784.4563 | 18.9 | 4 | y8 |
| Ribosomal protein L6 (YFP quantification) | APVVVPAGVDVK.light | 575.8452 | 685.3879 | 18.9 | 4 | y7 |

| | | | | | | |
|---|---|---|---|---|---|---|
| Ribosomal protein L6 (YFP quantification) | APVVVPAGVDVK.light | 575.8452 | 268.1656 | 18.9 | 4 | b3 |
| Ribosomal protein L6 (YFP quantification) | APVVVPAGVDVK.light | 575.8452 | 367.2340 | 18.9 | 4 | b4 |
| Ribosomal protein L6 (YFP quantification) | APVVVPAGVDVK.light | 575.8452 | 466.3024 | 18.9 | 4 | b5 |
| Ribosomal protein L6. QconCAT (YFP quantification) | APVVVPAGVDVK.heavy | 582.3259 | 893.4951 | 18.9 | 4 | y9 |
| Ribosomal protein L6. QconCAT (YFP quantification) | APVVVPAGVDVK.heavy | 582.3259 | 793.4296 | 18.9 | 4 | y8 |
| Ribosomal protein L6. QconCAT (YFP quantification) | APVVVPAGVDVK.heavy | 582.3259 | 693.3642 | 18.9 | 4 | y7 |
| Ribosomal protein L6. QconCAT (YFP quantification) | APVVVPAGVDVK.heavy | 582.3259 | 271.1567 | 18.9 | 4 | b3 |
| Ribosomal protein L6. QconCAT (YFP quantification) | APVVVPAGVDVK.heavy | 582.3259 | 371.2221 | 18.9 | 4 | b4 |
| Ribosomal protein L6. QconCAT (YFP quantification) | APVVVPAGVDVK.heavy | 582.3259 | 471.2876 | 18.9 | 4 | b5 |
| Ribosomal protein S1 (PSSA quantification) | GVVVAIDK.light | 400.7475 | 644.3978 | 13.4 | 4 | y6 |
| Ribosomal protein S1 (PSSA quantification) | GVVVAIDK.light | 400.7475 | 545.3293 | 13.4 | 4 | y5 |
| Ribosomal protein S1 (PSSA quantification) | GVVVAIDK.light | 400.7475 | 446.2609 | 13.4 | 4 | y4 |
| Ribosomal protein S1 (PSSA quantification) | GVVVAIDK.light | 400.7475 | 426.2711 | 13.4 | 4 | b5 |
| Ribosomal protein S1. QconCAT (PSSA quantification) | GVVVAIDK.heavy | 405.234 | 651.3770 | 13.4 | 4 | y6 |
| Ribosomal protein S1. QconCAT (PSSA quantification) | GVVVAIDK.heavy | 405.234 | 551.3115 | 13.4 | 4 | y5 |
| Ribosomal protein S1. QconCAT (PSSA quantification) | GVVVAIDK.heavy | 405.234 | 451.2461 | 13.4 | 4 | y4 |
| Ribosomal protein S1. QconCAT (PSSA quantification) | GVVVAIDK.heavy | 405.234 | 431.2563 | 13.4 | 4 | b5 |
| Ribosomal protein L1 (PSSA quantification) | VVGQLGQVLGPR.light | 611.8670 | 1024.5898 | 20 | 4 | y10 |
| Ribosomal protein L1 (PSSA quantification) | VVGQLGQVLGPR.light | 611.8670 | 839.5098 | 20 | 4 | y8 |
| Ribosomal protein L1 (PSSA quantification) | VVGQLGQVLGPR.light | 611.8670 | 726.4257 | 20 | 4 | y7 |
| Ribosomal protein L1 (PSSA quantification) | VVGQLGQVLGPR.light | 611.8670 | 442.2772 | 20 | 4 | y4 |
| Ribosomal protein L1 (PSSA quantification) | VVGQLGQVLGPR.light | 611.8670 | 329.1932 | 20 | 4 | y3 |
| Ribosomal protein L1 (PSSA quantification) | VVGQLGQVLGPR.heavy | 620.3418 | 1039.5453 | 20 | 4 | y10 |
| Ribosomal protein L1 (PSSA quantification) | VVGQLGQVLGPR.heavy | 620.3418 | 851.4742 | 20 | 4 | y8 |
| Ribosomal protein L1 (PSSA quantification) | VVGQLGQVLGPR.heavy | 620.3418 | 737.3931 | 20 | 4 | y7 |
| Ribosomal protein L1 (PSSA quantification) | VVGQLGQVLGPR.heavy | 620.3418 | 449.2565 | 20 | 4 | y4 |
| Ribosomal protein L1 (PSSA quantification) | VVGQLGQVLGPR.heavy | 620.3418 | 335.1754 | 20 | 4 | y3 |
| Ribosomal protein S4 (PSSA quantification) | AALELAEQR.light | 500.7747 | 745.3839 | 16.5 | 4 | y6 |
| Ribosomal protein S4 (PSSA quantification) | AALELAEQR.light | 500.7747 | 616.3413 | 16.5 | 4 | y5 |

| | | | | | | |
|---|---|---|---|---|---|---|
| Ribosomal protein S4 (PSSA quantification) | AALELAEQR.light | 500.7747 | 503.2572 | 16.5 | 4 | y4 |
| Ribosomal protein S4 (PSSA quantification) | AALELAEQR.light | 500.7747 | 432.2201 | 16.5 | 4 | y3 |
| Ribosomal protein S4 (PSSA quantification) | AALELAEQR.light | 500.7747 | 385.2082 | 16.5 | 4 | b4 |
| Ribosomal protein S4 (PSSA quantification) | AALELAEQR.heavy | 507.2555 | 755.3542 | 16.5 | 4 | y6 |
| Ribosomal protein S4 (PSSA quantification) | AALELAEQR.heavy | 507.2555 | 625.3146 | 16.5 | 4 | y5 |
| Ribosomal protein S4 (PSSA quantification) | AALELAEQR.heavy | 507.2555 | 511.2335 | 16.5 | 4 | y4 |
| Ribosomal protein S4 (PSSA quantification) | AALELAEQR.heavy | 507.2555 | 439.1994 | 16.5 | 4 | y3 |
| Ribosomal protein S4 (PSSA quantification) | AALELAEQR.heavy | 507.2555 | 389.1963 | 16.5 | 4 | b4 |

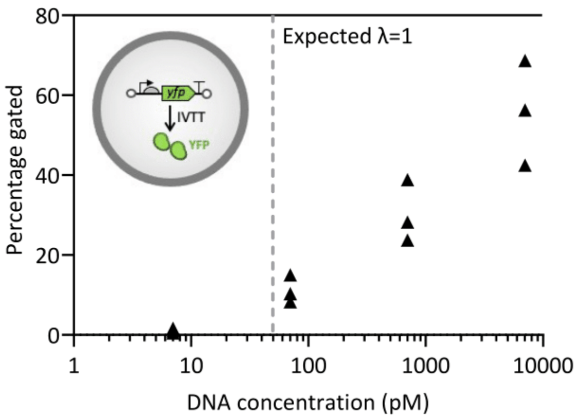


**Fig S1.** Effect of DNA concentration on YFP protein expression without gene amplification. Fluorescence from individual liposomes was analyzed by flow cytometry.

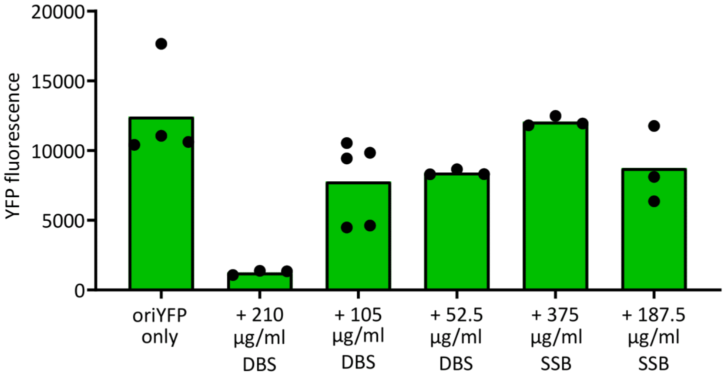


**Fig S2.** End-point YFP fluorescence measurements from ori-*yfp* bulk IVTT reactions. Protein expression can be inhibited under high DSB concentrations (210 µg/ml).

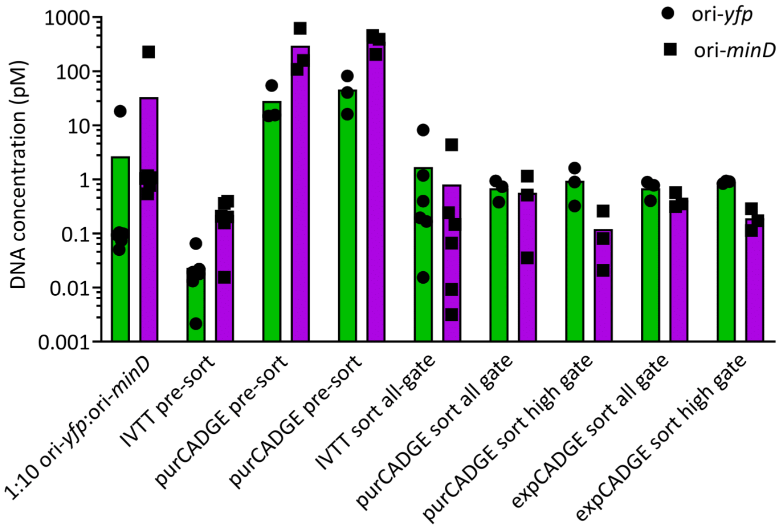


**Fig S3.** Individual qPCR data from Figure 4h. Enrichment of ori-*yfp* over ori-*minD*. Each symbol represents a biological repeat. 'IVTT' indicates a reaction without DNA replication.

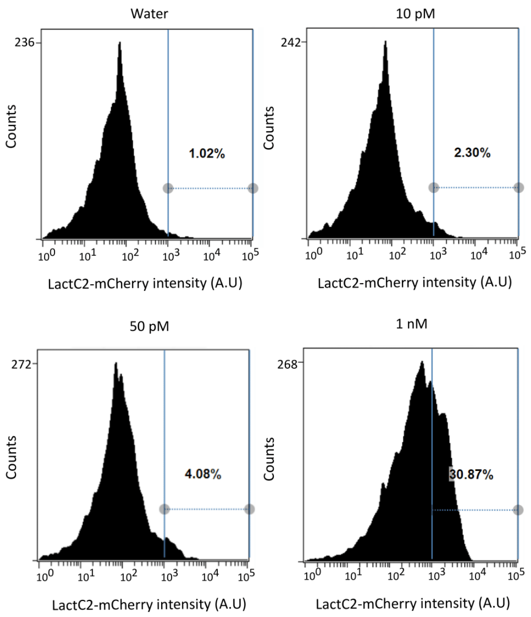


**Fig S4.** Raw FACS data of liposome samples with appended gating line as used in Figure 5c,d.

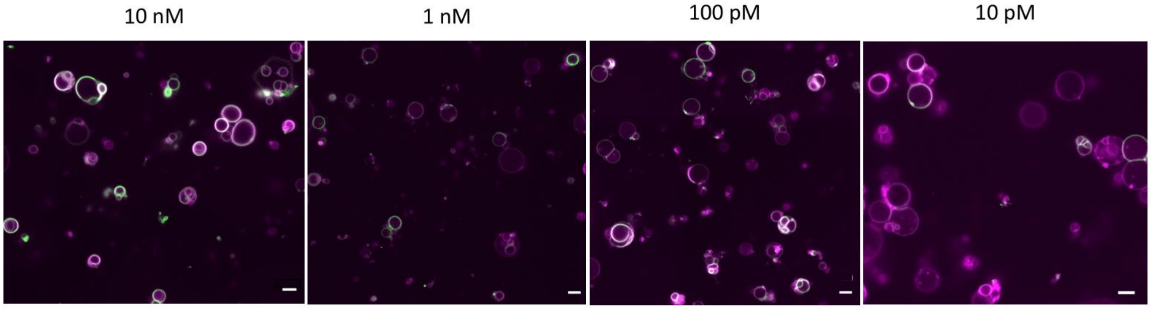


**Fig S5.** In-liposome expression of ori-*pssA* under different DNA concentrations (10 nM to 10 pM) without DNA replication. Liposome membrane dye (Texas-red) is colored in magenta and PS binding protein LactC2-eGFP is colored in green. Overlay of the two colors is displayed in white. Lowering the concentration of *pssA* gene reduces the number of liposomes with membrane-recruited LactC2-eGFP. Scale bars are 5 µm.

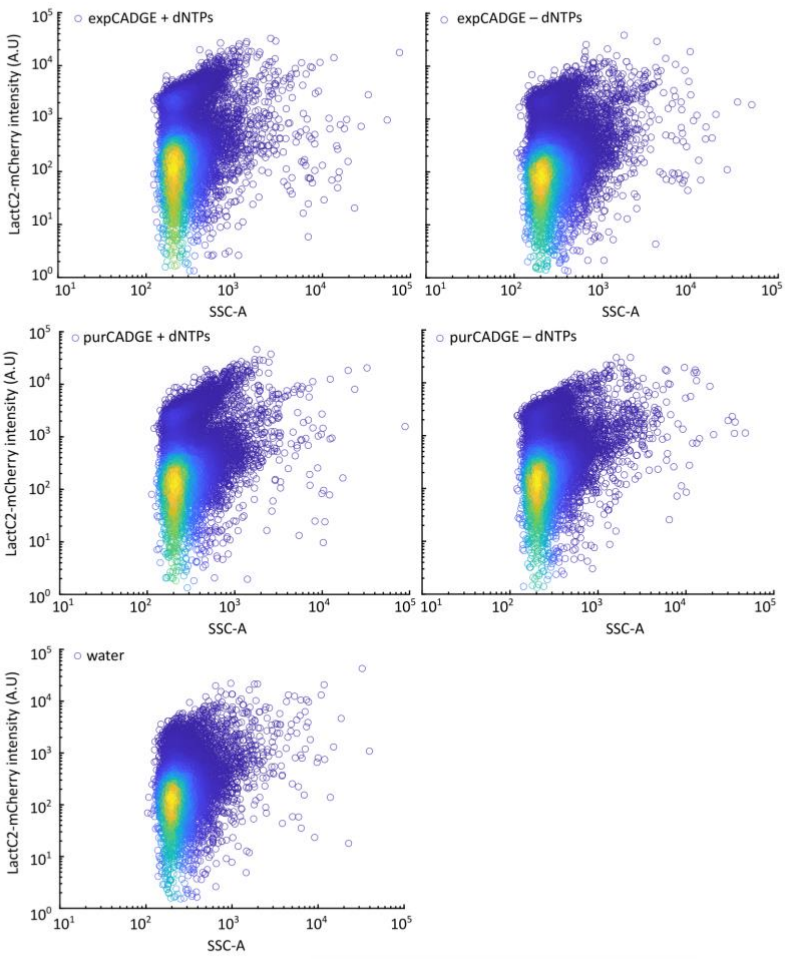


**Fig S6.** FACS data of liposome samples analyzed in Figure 5g.

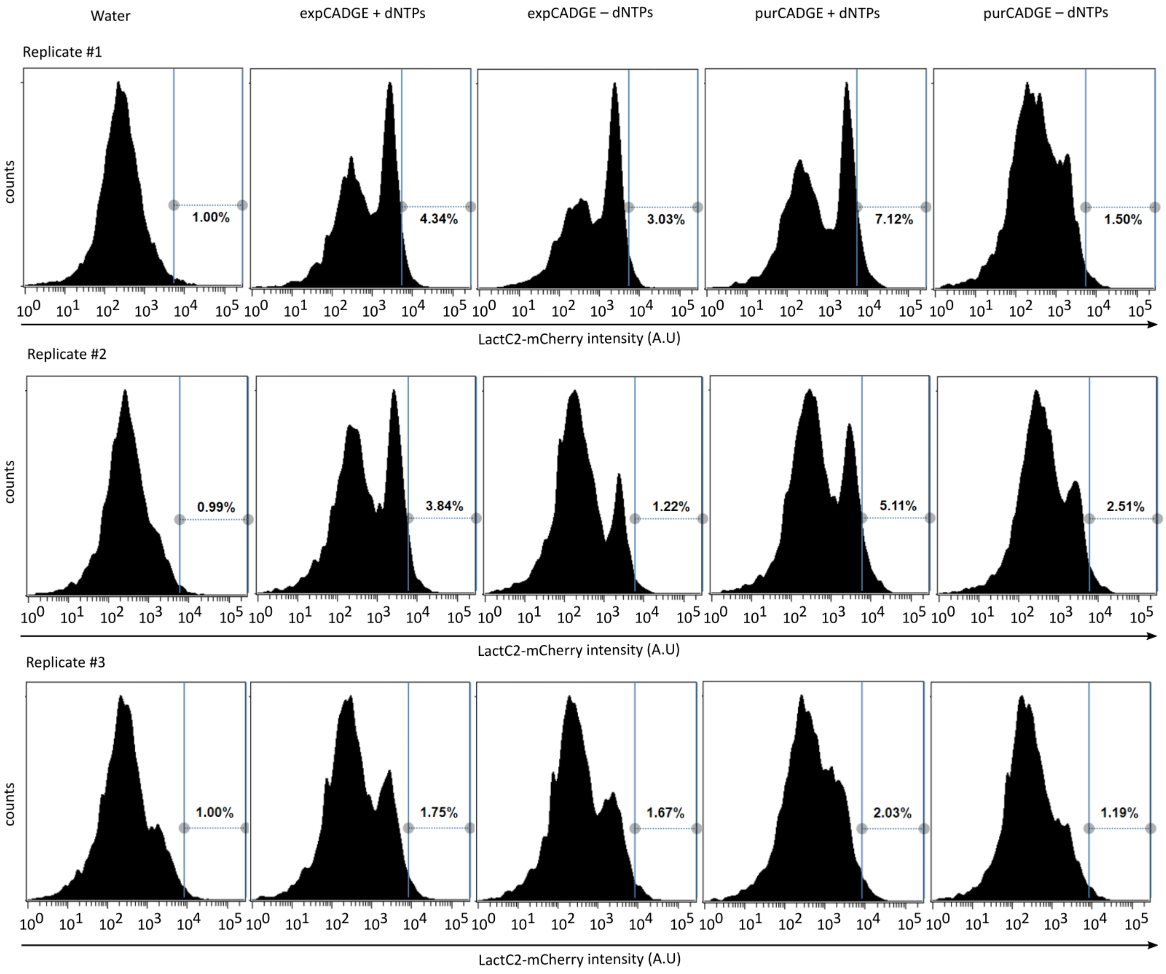


**Fig S7.** Liposomes FACS data and gating strategy for Figure 5h.

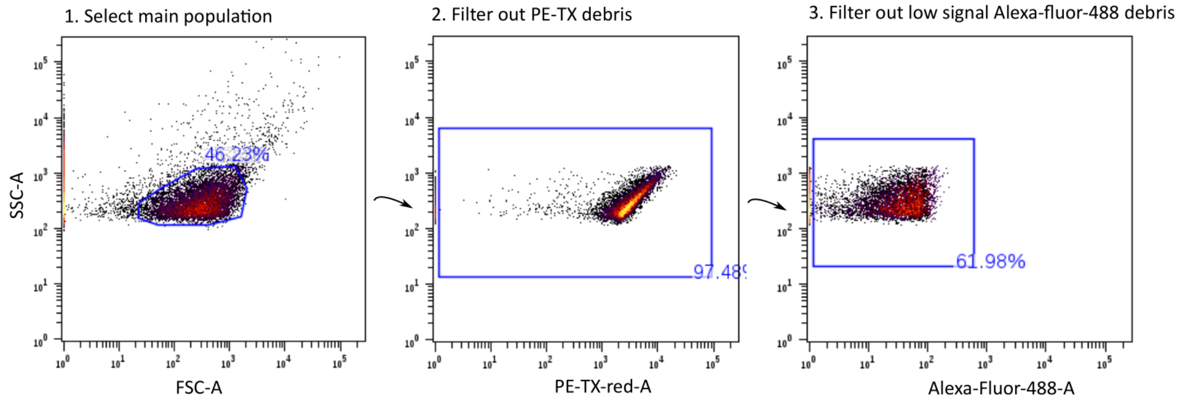


**Fig S8.** Data processing for FACS data. Application of the filtering gate to remove liposomal debris in PssA expressing-liposomes.
